# Whole genome sequencing reveals how plasticity and genetic differentiation underlie sympatric morphs of Arctic charr

**DOI:** 10.1101/2025.02.10.637319

**Authors:** Khrystyna Kurta, Mariano Olivera Fedi, Kendall Baker, Tom Barker, Leah Catchpole, Claudio Ciofi, Arianna Cocco, Genevieve Diedericks, Maria Angela Diroma, Kjetil Hindar, Alessio Iannucci, Naomi Irish, Vanda Knitlhoffer, Linda Laikre, Henrique G. Leitão, Sacha Lucchini, Seanna McTaggart, Arnar Pálsson, Mats E. Pettersson, Nils Ryman, Sigurður S. Snorrason, Hannes Svardal, David Swarbreck, Robert M. Waterhouse, Christopher Watkins, Han H. Xiao, Karim Gharbi, Zophonías O. Jónsson, Leif Andersson

## Abstract

Salmonids have a remarkable ability to form sympatric morphs after postglacial colonization of freshwater lakes. These morphs often exhibit differences in morphology, feeding, and spawning behaviour. Here we explore the genetics of morph differentiation by establishing a high-quality, annotated reference genome for the Arctic charr and use this as a resource for population genomic analysis of morphs from two Norwegian and two Icelandic lakes. The four lakes represent the spectrum of genetic differentiation between morphs from one lake with no genetic differentiation between morphs, implying phenotypic plasticity only, to two lakes with locus-specific genetic differentiation, implying incomplete reproductive isolation, and one lake with strong genome-wide divergence consistent with complete reproductive isolation. As many as 12 putative inversions ranging from 0.45 to 3.25 Mbp in size segregated among the four morphs present in one lake, Thingvallavatn, and these contributed significantly to the genetic differentiation among morphs. None of the putative inversions was found in any of the other lakes, but there were cases of partial haplotype sharing in similar morph contrasts in other lakes. The results are consistent with a highly polygenic basis of morph differentiation with limited genetic parallelism between lakes. The results support a model where morph differentiation is usually first established due to phenotypic plasticity that results in niche expansion and separation, followed by gradual development of reproductive isolation and locus-specific differentiation and eventually complete reproductive isolation and genome-wide divergence. A major explanation for salmonids ability to diversify into multiple sympatric morphs is likely the genome complexity caused by their ancient whole genome duplication that enhances evolvability.

## Introduction

One of the major challenges in evolutionary biology is to understand how species diversify into genetically distinct populations, subspecies and ultimately separate species ^1,2^. As the Arctic ice caps receded at the end of the last glacial epoch many species of salmonids colonized different freshwater systems across the Northern hemisphere. Salmonid species belonging to five different genera (*Coregonus*, *Oncorhynchus*, *Prosopium*, *Salmo,* and *Salvelinus*) are known for their ability to form genetically differentiated sympatric morphs that differ in morphology and/or feeding behavior (reviewed by Salisbury & Ruzzante, 2022). Considering that environmental factors are crucial drivers of sympatric differentiation ^4^, phenotypic plasticity has been regarded as a rapid response mechanism that could generate functional variation and foster adaptive evolution ^5,6^. Under conditions where the environmental pressures remain consistent over generations, these plastic responses can facilitate the establishment of genetically differentiated morphs ^7^. The presence of such sympatric morphs in freshwater systems across the Northern hemisphere provides an excellent opportunity to explore questions of fundamental importance in evolutionary biology, including the genetics of phenotypic differentiation, genetic parallelism, and mechanisms underlying reproductive isolation.

The Arctic charr (*Salvelinus alpinus*) has a Holarctic distribution and is found both as permanent freshwater resident and anadromous populations ^8,9^. It has colonized thousands of lake systems across the Holarctic region and in alpine regions of the boreal zone ^10^ and shows extensive phenotypic differentiation within and among lakes (see e.g. Adams & Maitland, 2007; Klemetsen, 2010). The co-existence of two or more discrete phenotypes (or morphs) of Arctic charr within a lake has repeatedly been reported (reviewed by ^3^). To name some well-known examples, genetically distinct small and large morphs of Arctic charr occur in lakes in Labrador, Canada ^13^ and dwarf, small, and large charr each occur in multiple lakes in northern Transbaikalia, Russia ^14^. In Iceland, genetically distinct sympatric morphs have been described in a number of lakes (^15,16^ and in one of these, Lake Thingvallavatn, four distinct morphs have been described ^17,18^. In Norway, several cases of sympatric populations have been described with two ^19,20^ to three genetically differentiated morphs ^21,22^. Given the rich data on morphology, diet, habitat use, spawning, life history, and genetic differentiation among Arctic charr morphs, they represent an excellent model for investigating the detailed genetic basis of sympatric divergence.

Genetic differentiation among sympatric morphs of Arctic charr, suggestive of at least partial reproductive isolation, is well documented ^7,15,23^. Furthermore, many studies suggest monophyly of sympatric morphs, as different morphs within the same lake exhibit closer genetic relatedness than similar morphs from different lakes ^13–15,24^. Importantly, the differentiation of Arctic charr into distinct morphs might occur via plastic responses during development, e.g. due to varying food resource utilization, without specific genetic factors involved ^25,26^. However, the absence of genetic contributions to phenotypic differentiation can only be concluded after whole genome sequencing (WGS) has been performed, because gene flow between sympatric populations can erase genetic differentiation at neutral loci. WGS has proven useful in many systems ^27^, and can unravel genetic changes that contribute to morph differentiation in Arctic charr populations and other salmonids. Furthermore, access to a high-quality reference genome is a prerequisite for high-resolution genomic, transcriptomic and epigenomic studies. However, omic studies in salmonids have been hampered by genome complexity due to an ancestral whole genome duplication that happened about 100 million years ago (Mya) ^28^. The varying degree of rediploidization and presence of highly similar paralogs make it challenging to establish high-quality genome assemblies and to accurately characterize many types of sequence variation. Previous studies on sympatric differentiation in salmonids have been based on restricted sets of genetic markers (microsatellites, AFLP - amplified fragment length polymorphisms, RADseq - restriction site associated DNA markers; Salisbury and Ruzzante 2022, but see Saha et al. 2022). To date, the available reference genomes that can be used for Arctic charr genomics is a genome assembly of a possible hybrid with a closely related species, the Northern Dolly Varden (*S. malma malma*: accession number GCA_002910315.2), a recently deposited scaffold-level assembly for a Canadian Arctic charr subspecies (*Salvelinus alpinus oquassa;* accession number GCA_036784965.1), and a genome assembly representing a selectively bred line of Arctic charr (*Salvelinus alpinus*; accession number GCA_045679555.1).

In the present study, we present a high-quality, chromosome-level scaffold genome for the Arctic charr with protein-coding gene annotation. We use this resource as a powerful tool for a population genomics project involving low-pass, whole genome sequencing of 283 individuals representing different morphs from two Icelandic and two Norwegian lakes where sympatric Arctic charr populations occur naturally and have been well characterized ^18,19^. The four lakes included in this study exhibit the range of genetic differentiation previously observed in salmonids ^3^, ranging from a lake with no/limited genetic differentiation between morphs, to one with strong genetic differentiation but noticeable gene-flow, and to a lake with strong genetic differentiation across the genome implying complete, or near complete, reproductive isolation. Combined across lakes, seven different morphs are included in the study: dwarf benthic (DB), large pelagic (LP), large generalist (LG), piscivorous (Pi), planktivorous (PL), large benthic (LB), and small benthic (SB). This study is the first attempt to explore the genetics of sympatric differentiation in Arctic charr based on whole genome sequencing. By reanalyzing the original samples from the pioneering genetic studies in Norwegian charr ^19^ alongside new samples from ongoing research in Iceland utilizing ddRAD-sequencing ^29^; see Supplementary Table 1), this work provides a comprehensive and updated genome-wide analysis of Arctic charr morph differentiation.

## Results

### Genome assembly overview

We assembled the genome of a male large benthic Arctic charr sampled from Lake Thingvallavatn in Iceland using 8.7M Pacific Biosciences high-fidelity (HiFi) reads with a mean length of 18 kb, generating approximately 52-fold coverage of the haploid genome (Table 1). Primary assembly contigs were scaffolded with Hi-C chromosome conformation data from 386M Illumina short reads, providing approximately 38-fold coverage. The final, manually curated assembly has a total length of 2.99 Gb in 5,797 sequence scaffolds with a scaffold N50 of 53.2 Mb (Table 2). With the exception of BUSCO metrics showing an excess of duplicated genes consistent with tetraploid ancestry, assembly metrics exceeded Earth BioGenome Project (EBP, https://www.earthbiogenome.org/report-on-assembly-standards) standards ^30^ and were broadly similar to those of published assemblies derived from other salmonid species using equivalent data types, including HiC (e.g. ^30–33^. We also found extensive synteny between our assembly and the high-density linkage map of Canadian Arctic charr ^34^ using GBS reads as anchors between scaffolds and linkage groups (Supplementary Figure 1). We annotated protein-coding genes in the assembly using short-read RNA-Seq alignments and transcript assemblies from Pacific Biosciences Iso-Seq reads, including 467.8M RNA-Seq and 7.3M Iso-Seq reads from seven adult tissues and two development stages (Table 1), as well as cross-species alignment of protein sequences. In total, we identified 47,703 protein-coding genes and 179,230 transposable elements with high-confidence.

**Table 1.** Genome assembly data summary.

| Technology | ENA accession | Read count | Base count (Gb) |
| --- | --- | --- | --- |
| HiFi PacBio Sequel IIe | PRJEB76174 | 8,677,046 | 157.07 |
| Hi-C Illumina NovaSeq 6000 | PRJEB76174 | 386,000,000 | 115.0 |
| RNA Illumina NovaSeq 6000 | PRJEB76174 | 467,823,625 | 140.3 |
| IsoSeq PacBio Sequel IIe | PRJEB76174 | 7,273,335 | 23.05 |

**Table 2.** Genome assembly and annotation statistics.

| Assembly name | fSalAlp1_hap1 |
| --- | --- |
| Span (bp) | 2,999,192,943 |
| Number of scaffolds | 5,797 |
| Scaffold N50 length (bp) | 53,120,343 |
| Consensus quality (QV) | 56.34 |
| *k*-mer completeness | 95.28% |
| BUSCO | C:97.2%, S:51.3%, D:45.9% |
| Number of protein-coding genes | 47,703 |
| Number of protein-coding transcripts | 128,882 |
| Number of transposable element genes | 179,230 |
| Number of transposable element transcripts | 13,030 |

### Genome-wide differentiation and population structure

Low-pass, whole genome resequencing was performed on 283 Arctic charr from two Icelandic and two Norwegian lakes (Fig. 1; Supplementary Table 1), each harboring two or four different morphs. Sequence coverage per individual was on average 2.1±0.9X (range: 0.22-6.11X). Inference of population structure was done using genotype likelihoods from the ANGSD software (Korneliussen et al., 2014). As expected from the geographic distances between these lakes and the topography of their out-flowing rivers, we found strong genetic differentiation between lakes (Fig. 2a-c). Furthermore, and consistent with earlier studies ^19,29,35,36^, different morphs from the same lake clustered tightly along the first three principal component (PC) axes. Admixture analysis with PCAngsd gave the highest support at *K* = 5 and, as in the case of the PC analysis, revealed genetic separation by lakes, but also two distinct clusters in Lake Sirdalsvatnet (Fig. 2d). Consistently, the neighbor-joining tree showed four clusters separating the four lakes with an additional split in Sirdalsvatnet highlighting strong genetic divergence between the dwarf benthic and large pelagic morphs in this lake (Fig. 2e). Pairwise *Fst* values between population pairs (i.e. lakes) ranged from 0.21±0.15 to 0.28±0.18 (Fig. 2f), further indicating substantial genetic differentiation among lakes.

**Fig. 1.**
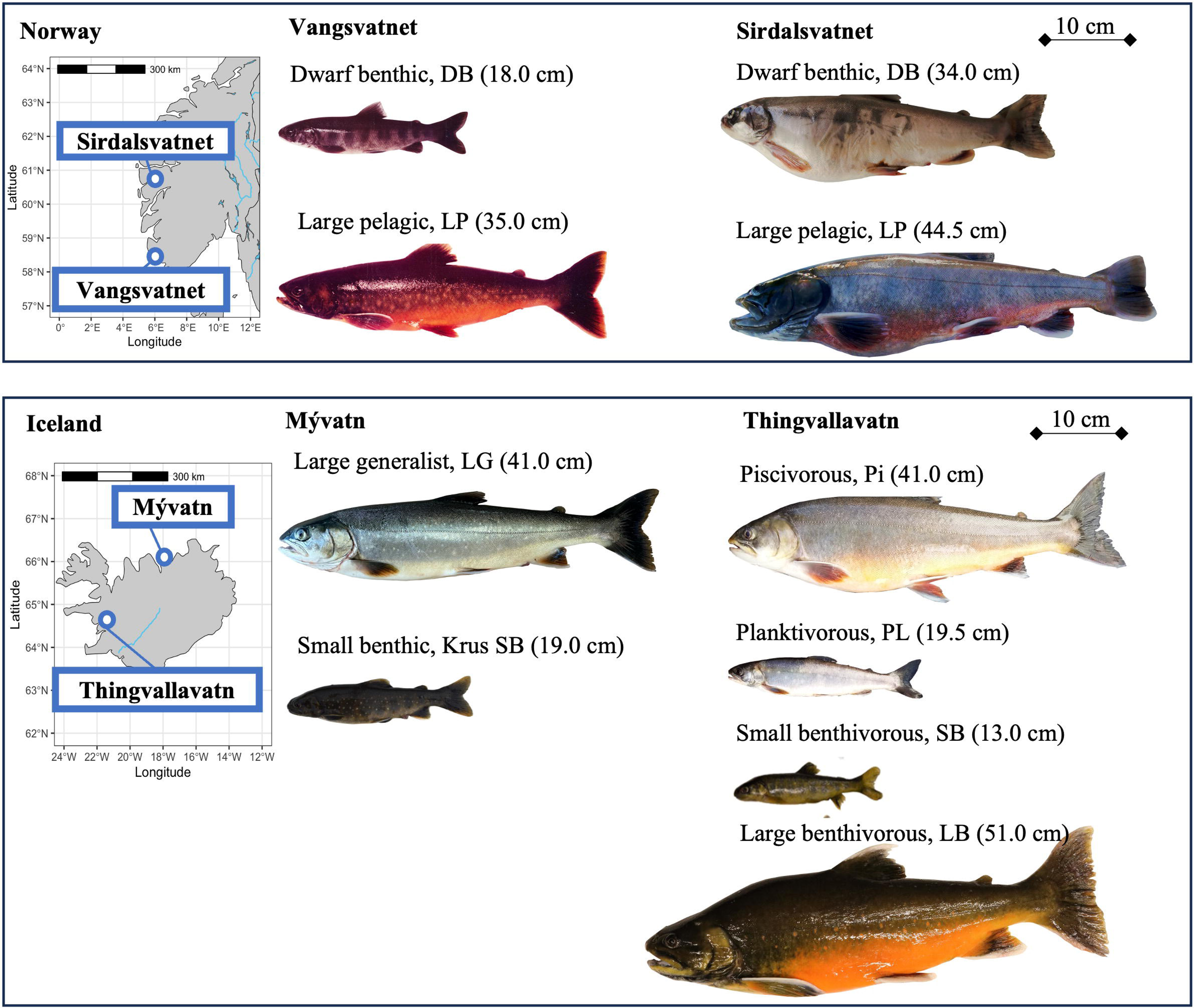
Geographic location and phenotypic differentiation of Arctic charr (*Salvelinus alpinus*) morphs from Norway (Vangsvatnet and Sirdalsvatnet) and Iceland (Mývatn and Thingvallavatn). Fork lengths of the fish are shown in brackets next to the morph name. The reference genome generated in this study came from the large benthivorous charr (LB-charr) shown in the bottom right corner. Photo credits: Tormod A. Schei (Vangsvatnet), Ragnvald Andersen (Sirdalsvatnet), Gylfi Yngvason and Árni Einarsson (LG, Mývatn); Sigurður S. Snorrason and Zophonías O. Jónsson (SB, Mývatn); Sigurður S. Snorrason, Zophonías O. Jónsson, and Arnar Palsoon (Thingvallavatn).

**Fig. 2.**
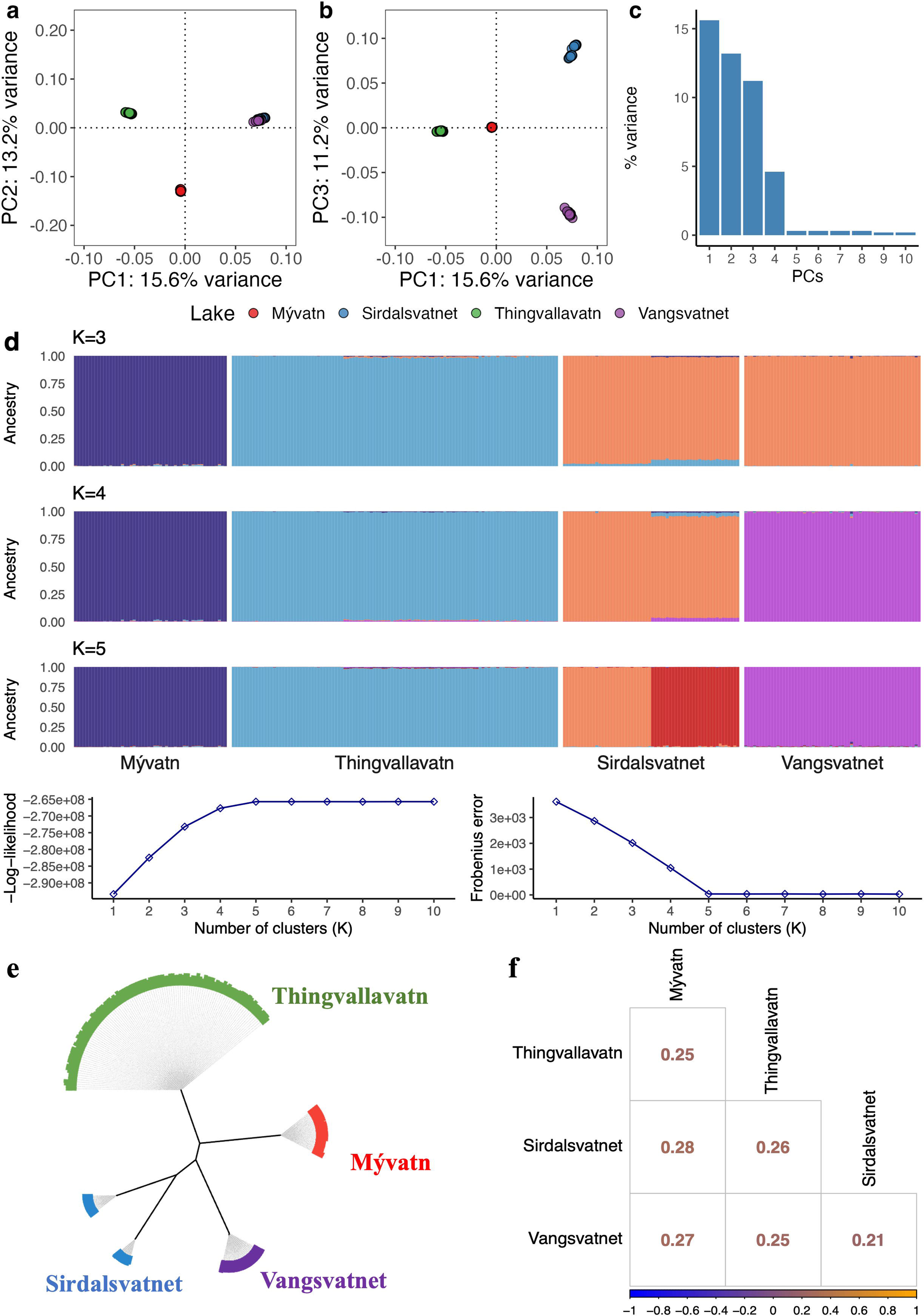
Genetic differentiation in Arctic charr among lakes and morphs. PCA, Admixture, and Neighbor-joining tree analyses generated in PCAngsd based on a downsampled list of ∼1 million SNPs (MAF >0.05). Scores of individuals along principal components **(a)** PC1 and PC2 and **(b)** PC1 and PC3), and **(c)** proportion of explained variance across ten principal components. (**d)** Ancestry proportions with -Log-likelihoods and Frobenius error for 1–10 clusters. **(e)** Unrooted Neighbor-joining tree and **(f)** pairwise Fst values illustrating genetic differentiation between lakes.

### Genetic differentiation and population structure between morphs within each lake

We applied principal component analysis and estimated admixture proportions to evaluate genetic differentiation among sympatric morphs within lakes and performed genome-wide contrasts (GWC) to detect specific loci contributing to genetic differentiation between morphs. First, we compared dwarf benthic and large pelagic morphs in the Norwegian lakes Sirdalsvatnet and Vangsvatnet and found high genome-wide divergence between the morphs from Sirdalsvatnet (Fig. 3a,b). Admixture analysis showed the highest support at *K* = 2, separating the two morphs in Sirdalsvatnet (Fig. 3b). The average *Fst* value between the two morphs in Sirdalsvatnet was 0.13±0.14 (mean±s.d). In sharp contrast, the corresponding morphs (dwarf benthic and large pelagic) in Vangsvatnet were not differentiated genetically (Fig. 3c,d), and average *Fst* was as low as 0.01±0.00 (mean±s.d). The GWC detected genome-wide differentiation between the Sirdalsvatnet morphs (Supplementary Fig. 3a, Supplementary Table 2), consistent with strong reproductive isolation and genetic differentiation at both neutral and adaptive loci. Genetic differentiation may have occurred sympatrically or alternatively the two populations may have populated the lake separately. As expected, no significant differentiation was found in a GWC between the two morphs in Lake Vangsvatnet (Supplementary Fig. 3b). This could indicate that the two morphs in Vangsvatnet are not reproductively isolated.

**Fig. 3.**
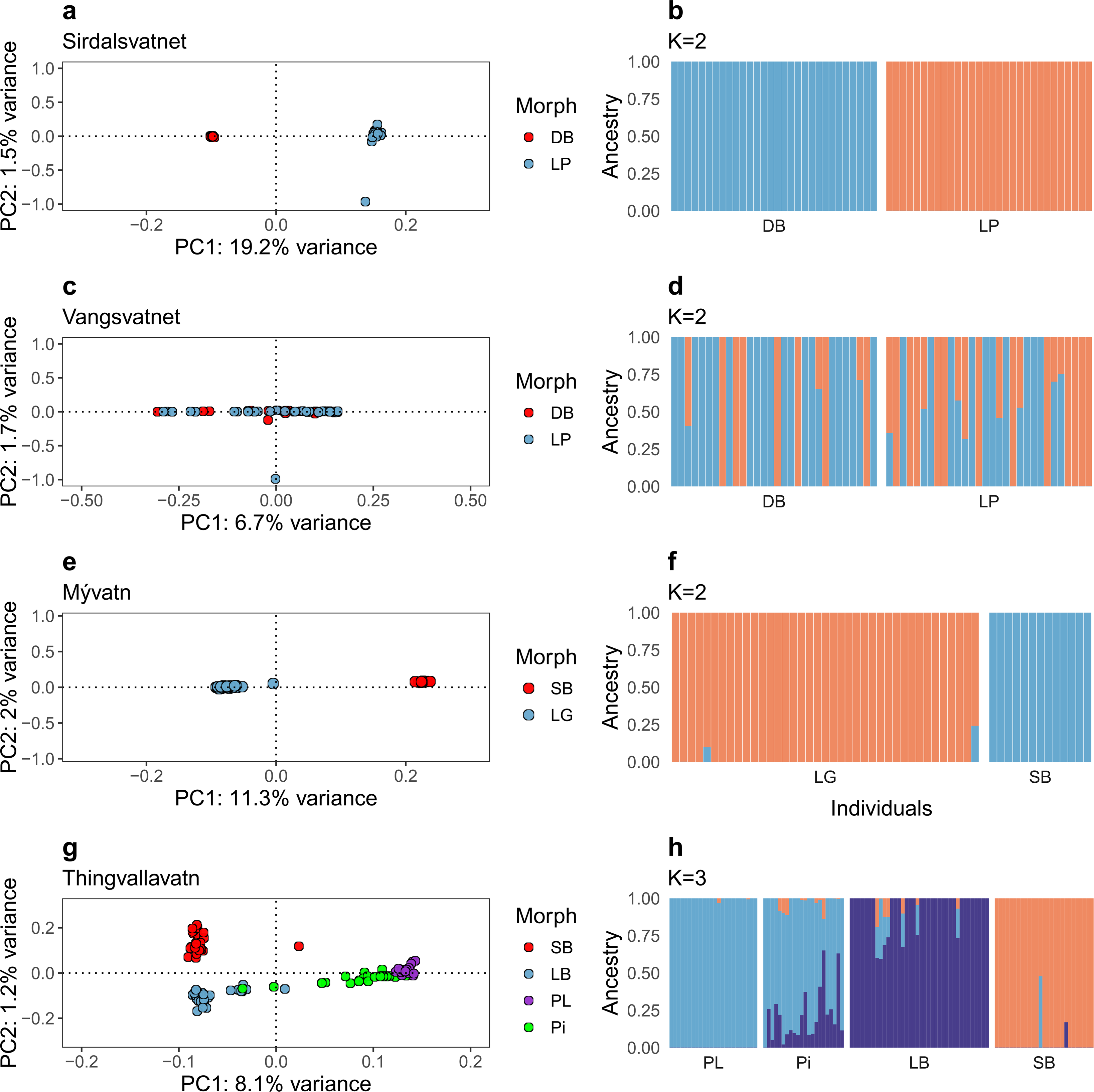
Genetic differentiation among Arctic charr morphs in two Norwegian and two Icelandic lakes. Scores of individuals along PC1 and PC2 with the genetic variance explained from PCA (left), and ancestry proportions (right) for each lake. PC scores were generated using PCAngsd, and individual ancestry proportions were estimated with NGSadmix based on a downsampled list of SNPs and MAF > 0.05 in every lake. The numbers of SNPs used per population: **(a, b)** Sirdalsvatnet (0.73 million SNPs), (**c, d)** Vangsvatnet (0.65 million SNPs), **(e, f)** Mývatn (0.51 million SNPs), and **(g, h)** Thingvallavatn (0.58 million SNPs). Colors on PCA plots indicating the sympatric morphs of Arctic charr for Sirdalsvatnet and Vangsvatnet: Dwarf benthic (DB) and Large pelagic (LP); for Mývatn, Large generalist (LG) and Small benthic (Krús, SB), and Thingvallavatn, Piscivorous (Pi), Planktivorous (PL), Large benthivorous (LB) and Small benthivorous (SB).

In Lake Mývatn, we compared fish classified as small benthic and large generalist. PC analysis showed two distinct genetic groups separated along the PC1 axis that explained 11.3% of variance (Fig. 3e). Additionally, admixture analysis suggested the presence of *K* = 2 distinct genetic backgrounds with minor evidence of shared genetic variation, as indicated by two LG individuals exhibiting ∼10-24% admixture from the SB group (Fig. 3f). The average *Fst*±s.d. (0.05±0.03) indicates a moderate level of genetic differentiation between morphs in Mývatn. A GWC contrasting these genetically classified morphs detected single nucleotide polymorphisms (SNPs) exceeding Bonferroni-adjusted significance threshold across 20 scaffolds (Supplementary Fig. 3c, Supplementary Table 2).

Lake Thingvallavatn harbours four morphs and the first PC (8.1% of variance) separated the benthic morphs from the planktivorous morph and the majority of piscivorous individuals, while PC2 (1.2% of variance) separated the small and large benthivorous morphs (Fig. 3g). The result confirms previously published findings based on allozyme loci ^37^ and microsatellites ^36^. As previously reported ^7^, most piscivorous individuals grouped with or close to the planktivorous charr cluster, however several individuals fell in between the large benthivorous and piscivorous/planktivorous clusters. Estimated admixture proportions with PCAngsd suggested *K* = 2 genetic backgrounds, likely corresponding to the broad separation of benthivorous and piscivorous/planktivorous morphs, as revealed by PC1. However, NGSadmix detects an additional layer of genetic structure (*K* = 3, Supplementary Fig. 4a, b), consistent with the separation of the small and large benthivorous morphs (captured by PC2). with evidence of shared genetic variation between the four morphs (Fig. 3h). Notably, the *Fst* values (Supplementary Fig. 4c) for the benthivorous versus planktivorous morph pairs suggested moderate differentiation (*Fst* in the range 0.03±0.03 to 0.05±0.07), while the piscivorous/planktivorous contrast exhibited minimal differentiation (*Fst* = 0.01±0.01). Consistent with these data, the GWC detected genome-wide significant regions of genetic differentiation at many loci across the genome between all morph pairs from Thingvallavatn (Supplementary Table 2, Supplementary Fig. 3d-i) except for the piscivorous/planktivorous contrast, where a single significant SNP was found on scaffold 15 (Supplementary Fig.3 i).

### Genetic differentiation at putative inversions present in Lake Thingvallavatn

We next investigated the genomic regions that differentiate the two benthivorous morphs (LB and SB) in Lake Thingvallavatn. Highly differentiated SNPs above the Bonferroni threshold (*P* <1×10^−10^) were detected across 33 scaffolds and unplaced scaffolds (Fig. 4a). Notably, four of the major outlier loci involved large genomic regions spanning 0.88 Mb on scaffold 4, 0.45 Mb on scaffold 5, 0.81 Mb on scaffold 9, and 0.75 Mb on scaffold 17 (Fig. 4b-e). A closer look at these loci revealed a block-like pattern with many SNPs showing strong genetic differentiation and with sharp drops in differentiation at each end of the block, a pattern consistent with the presence of inversions (see e.g., Han et al., 2020). The linkage disequilibrium (LD) among SNPs across all four inversions was strong, particularly for scaffolds 4, 5 and 17 inversions (Fig. 4b-e). These putative inversions contain altogether 64 genes: 30 on scaffold 4, one on scaffold 5, one on scaffold 9, and 32 on scaffold 17 (Supplementary Table 9). Notably, some of the detected genes within these inversions were previously identified as genetically differentiated based on transcriptome sequencing, including *WEE1*, *GAS1*, and *DEN5A* ^7^. *WEE1* is a critical regulator of cell cycle progression and mitosis ^38^. *GAS1* is associated with embryonic cranial skeleton development and palate formation ^39^, while *DEN5A* is involved in the negative regulation of neurite outgrowth ^40^. A summary of the most differentiated SNPs (*P* <1×10^−15^) for each region, including their nearest gene, and relative position to genes (e.g., nonsynonymous, synonymous, upstream, downstream, or intergenic) as determined by snpEff are presented in Supplementary Table 12. We calculated nucleotide diversity (θ) for each haplotype within the four putative inversions. First, we determined the genotype at each locus (Supplementary Table 3) and calculated nucleotide diversity for individuals homozygous for either the small or the large benthivorous-associated inversion haplotype (Fig. 4f). While the genome-wide nucleotide diversity was very similar in the two morphs (dwarf: 0.15%; large benthic: 0.14%), the average θ values within the putative inversions were significantly higher (Wilcoxon test, *P* < 0.01) in the small (from 0.08% to 0.28%) than in the large benthivorous morph (from 0.06% to 0.14%) at all four loci (Fig. 4f, Supplementary Table 4). We further explored the patterns of genotypes over these putative inversions, using only diagnostic markers (*P* <1×10^−15^) among morphs from Thingvallavatn and the other lakes (Supplementary Fig. 5). This analysis illustrates the striking genetic differentiation between small and large benthivorous morphs of Thingvallavatn at these loci. Furthermore, the large benthivorous haplotypes at scaffolds 4, 5 and 17 were strikingly similar to the haplotypes dominating among the pelagic morphs in the lake. We also explored the possibility that these putative inversions might segregate in any of the other three lakes included in this study, but we found no evidence for a similar pattern involving the entire regions (Supplementary Fig. 5). However, there was an interesting pattern at two relatively large fragments (61,387,699-61,393,705 bp; and 61,5015,90-61,615,879 bp) within the putative inversion region on scaffold 9: 61.3-62.1 Mb, in which the small benthic morphs from Thingvallavatn (Iceland) and Sirdalsvatnet (Norway) shared haplotypes (Supplementary Fig. 5), whereas the planktivorous morph from Sirdalsvatnet shared the haplotypes of the large benthivorous morph from Thingvallavatn. This haplotype sharing represents ancestral polymorphisms that may have contributed to morph differentiation in many lake systems and alleles at these ancestral polymorphisms became locked within the putative inversion segregating in Thingvallavatn. Another interesting pattern was noted for scaffold 17 involving the same comparison of morphs from Thingvallavatn and Sirdalsvatnet (Supplementary Fig. 5). However, in this case the haplotype present in the planktivorous morph in the latter lake is characterized by strong genetic differentiation at dispersed SNPs across the region, suggesting that recombination or a selective sweep has occurred.

**Fig. 4.**
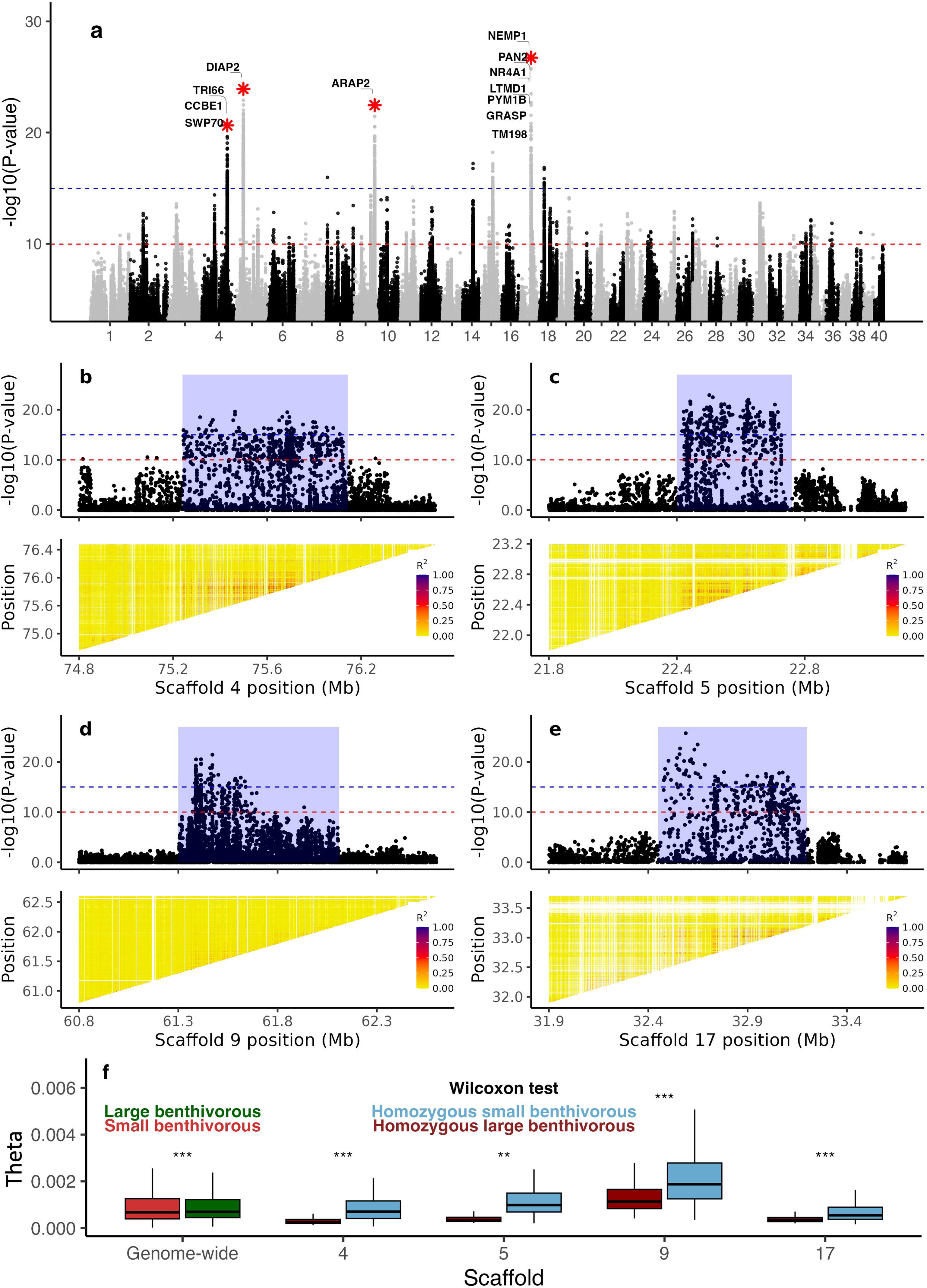
Genetic differentiation between small and large benthivorous morphs of Arctic charr in Thingvallavatn. (**a)** Genome scan based on estimated allele frequencies for individual SNPs. Gene annotations are shown for the top 0.001% of significant SNPs at putative inversions marked by stars. **(b-e)** Zoom-in profile and linkage disequilibrium represented as R^2^ among genotypes within the four putative inversion regions in **(a)**, including the following genomic regions **(b)** Scaffold 4: 75.25-76.13 Mb, **(c)** Scaffold 5: 22.30-22.75 Mb, **(d)** Scaffold 9: 61.30-62.11 Mb, **(e)** Scaffold 17: 32.45-33.20 Mb. The horizontal red and blue lines indicate Bonferroni corrected significance thresholds with significance levels *α =* 10^−3^ and *α =* 10^−8^, respectively, after taking into account that 10^7^ SNPs were used. **(f)** Wilcoxon rank test of pairwise theta (nucleotide diversity) distributions across the genome and within putative inversion regions.

We next pooled all benthivorous (SB and LB) individuals as one group and compared allele frequencies SNP-by-SNP with data consisting of all pelagic (Pi and PL) individuals from Thingvallavatn. The analysis detected extremely differentiated SNPs (*P* <1×10^−15^) across 37 scaffolds (Fig. 5a). Moreover, similar to the observed contrast between dwarf and large benthivorous morphs, eight prominent outlier loci on six scaffolds exhibited a pattern suggesting putative inversions, spanning large regions of 2.3 and 2.7 Mb on scaffold 1, 2.3 and 3.25 Mb on scaffold 3, 0.78 Mb on scaffold 8, 2.4 Mb on scaffold 9, 0.54 Mb on scaffold 14, and 0.76 Mb on scaffold 40 (Fig. 5b-g). There is strong LD among alleles within the eight inversions (Fig. 5b-g). These regions harbor a total of 211 genes with 28 genes on scaffold 1:16.3-18.6 Mb, 42 on scaffold 1:19.5-22.2 Mb, 27 on scaffold 3: 33.5 - 35.8 Mb, 61 on scaffold 3: 37.3 - 40.6 Mb, 11 on scaffold 8, 23 on scaffold 9, 5 on scaffold 14, and 14 on scaffold 40 (Supplementary Table 10). A previous transcriptome study revealed SNPs in *COX11* and *LRRC1* located within the putative inversion that showed strong genetic differentiation between morphs consistent with the present study ^7^. Furthermore, *AP2* and *KDM5C* also located within the putative inversion regions in the benthic/pelagic contrast were found to be differentially expressed between benthic and limnetic charr ^7^. Additionally, *TIMP2*, a gene primarily involved in regulating peripheral extracellular matrix remodeling ^41^ showed a 3.8-fold higher expression in the developing head in charr embryos of benthic morphs compared with pelagic morphs from Lake Thingvallavatn ^42^. Furthermore, the aforementioned *AP2* gene, was previously shown to be associated with craniofacial development in zebrafish ^43^, which highlights its potential role in formation of morph-specific craniofacial traits in charr. Supplementary Table 12 summarizes the most differentiated SNPs (*P* <1×10^−15^) for each region and their effects in relation to the closest genes.

**Fig. 5.**
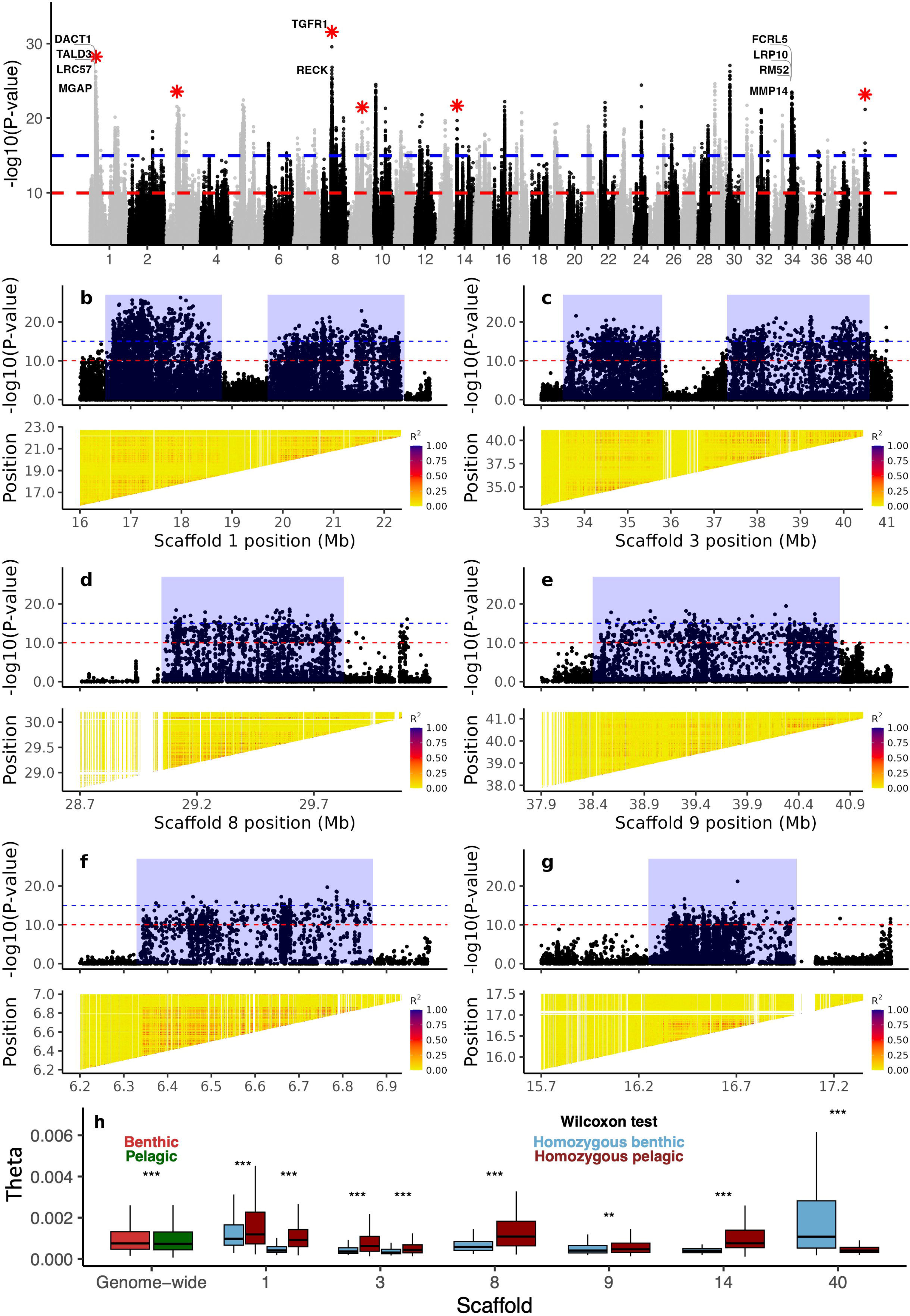
Genetic differentiation between benthic (large and small benthivorous) and pelagic (piscivorous and planktivorous) Arctic charr morphs in Thingvallavatn. **(a)** Genome scan based on estimated allele frequencies for individual SNPs. Gene annotations are shown for the top 0.001% of significant SNPs at putative inversions marked by stars and at scaffold 34. **(b-g)** Zoom-in profile and linkage disequilibrium represented as R^2^ among genotypes within the eight putative inversion regions in **(a),** including **(b)** Scaffold 1: 16.30-18.60 Mb and 19.50-22.20 Mb, **(c)** Scaffold 3: 33.50-35.80 Mb and 37.35-40.60 Mb, **(d)** Scaffold 8: 29.05-29.83 Mb, **(e)** Scaffold 9: 38.40-40.80 Mb, **(f)** Scaffold 14: 6.33-6.87 Mb, **(g)** Scaffold 40: 16.25-17.01 Mb. The horizontal red and blue lines indicate Bonferroni corrected significance thresholds with significance levels *α =* 10^−3^ and *α =* 10^−8^, respectively, after taking into account that 10^7^ SNPs were used. **(h)** Wilcoxon rank test of pairwise theta (nucleotide diversity) distributions across the genome and within putative inversion regions.

We estimated nucleotide diversity per haplotype at each inversion locus using individuals homozygous for the benthic and pelagic inversion haplotypes (Supplementary Tables 5 and 6, Fig. 5h). The analysis revealed consistent and significantly lower nucleotide diversities (Wilcoxon test, *P* < 0.01) among benthic inversion haplotypes at seven out of eight loci despite very similar genome-wide nucleotide diversity in the two morph groups (Fig. 5h). The pattern of genotype distributions at the eight putative inversion loci was further explored using diagnostic markers (*P* <1×10^−15^) among benthic versus pelagic morphs from Thingvallavatn as well as other lakes (Supplementary Fig. 6). This analysis showed that none of the putative inversions detected in Thingvallavatn is shared intact with the other lakes, suggesting that if these are true inversions they do not have a wide geographic distribution. However, there were multiple examples of haplotype sharing for parts of the putative inversions for which the pattern was consistent across morphs and lake systems. Patterns of haplotype sharing were observed between Thingvallavatn and Mývatn morphs along three distinct regions (16,453,643-16,910,602 bp; 17,026,128-170,573,48 bp; 17,464,035-17,653,321 bp) of the putative inversion on scaffold 1 (16.3-18.6 Mb). Additionally, similar patterns of differentiation were evident between Thingvallavatn and Sirdalsvatnet morphs along parts of the putative inversions 16.3-18.6 Mb (17,464,035-17,653,321 bp) and 19.5-22.2 Mb (20,429,418-20,718,588 bp) on scaffold 1, and within putative inversion 38.4-40.8 Mb (39,292,876-39,384,683 bp) on scaffold 9. An exception to the rule that the small and large benthivorous morphs shared haplotypes at these putative inversions occurs at scaffold 8, where the large benthivorous morph share haplotypes with the pelagic and piscivorous morphs (Supplementary Figure 6).

### Genetic differentiation and population structure between morphs across lakes

The analysis of the putative inversion regions suggested the existence of some haplotype sharing between the same type of morphs across lakes. We therefore extended this analysis to the remaining part of the genome outside the putative inversions. Thingvallavatn was selected as the starting point for our analyses due to the clear signals of genetic differentiation among morphs in this lake.

We screened for the presence of shared patterns of differentiation between morphs from different lakes using the most differentiated genomic regions (Bonferroni-corrected significance threshold with *α =* 10^−8^) obtained in the dwarf/large benthivorous (Fig. 4a) and benthic/pelagic morph contrasts (Fig. 5a) from Thingvallavatn. Only two genomic regions outside the putative inversions showed strong differentiation between dwarf and large benthivorous morphs in Thingvallavatn, but no similar patterns were found across lakes (Supplementary Fig. 7).

On the other hand, there was a notable example of shared patterns of genetic differentiation between lakes when analyzing regions that distinguish benthic and pelagic morphs in Thingvallavatn, in particular for scaffold 34 and the comparison between Thingvallavatn and Mývatn. The heatmap of allele frequency distributions across lakes (Supplementary Fig. 8) along with the corresponding predicted genotypes for individual fish (Supplementary Fig. 9a) showed that the small benthic morphs from Mývatn carried a haplotype for the region on scaffold 34: 18.36-18.45 Mb closely related to the one dominating among benthic morphs from Thingvallavatn. There was a moderate to strong LD (*R*^2^ range: 0.5 to 1.0) among genotypes within both lakes for this region (Supplementary Fig. 9b). This genomic region contains seven genes, *LRP10*, *MMP14*, *ACINU*, *AJUBA, CD244, FCRL5,* and *RM52* with the currently known functions: LRP10 involved in lipid metabolism ^44^; heterochrony in MMP14 expression along with other genes was associated with terminal/benthic mouth phenotype in sucker fish ^45^; *ACINU* plays a role in splicing regulation as well as in other cellular pathways ^46^; AJUBA may be involved in regulating cell growth and differentiation decisions during early development ^47^; *CD244* is an immunoregulatory receptor found on a variety of immune cells ^48^; *FCRL5* is also involved in immune system regulation ^49^; and *RM52* plays a critical role in the maintenance of genome integrity under oxidative stress conditions ^50^. A summary of the annotated genes and the most highly differentiated SNPs (*P* <1×10^−15^) can be found in Supplementary Tables 11 and 12, respectively.

To validate the presence of shared haplotypes, we carried out principal component analysis for individuals homozygous for the benthic (59 individuals) and pelagic (71 individuals) haplotypes from Thingvallavatn and Mývatn (Supplementary Table 8). The analysis was based on a set of markers (5,000 SNPs) located within the region 18.36-18.45 Mb of scaffold 34. The PCA explained a substantial proportion of variance (PC1: 33.3% and PC2: 22.8%) and illustrated distinct clustering of homozygous benthic and pelagic inversion haplotype samples from the two lakes (Supplementary Fig. 9c). Notably, PC2 further differentiated between benthic and pelagic individuals from Thingvallavatn and Mývatn, highlighting their genetic divergence despite observed haplotype sharing. To further explore this genetic structure, we constructed two Neighbor-joining phylogenetic trees of individuals being homozygous for benthic and pelagic haplotypes from the two lakes, both within and outside the shared region on scaffold 34 (Supplementary Fig. 8d, e). The clustering within the shared region (Supplementary Fig. 9d) was consistent with PCA results, and followed the classification of morphs across lakes, whereas the results for the region outside the shared haplotype showed a clustering based on lake as for the rest of the genome (Supplementary Fig. 9e). The branch lengths separating the clusters of individuals with benthic haplotypes from different lakes or between pelagic haplotypes from different lakes were shorter than between haplotypes within lakes (Supplementary Fig. 8d). Pairwise *Fst* values among two benthic groups (0.12) and within two pelagic groups (0.21) were lower compared to those estimated across groups (Mývatn: *Fst* = 0.81; Thingvallavatn: *Fst* = 0.81). This region on scaffold 34 constitutes another possible example of a shared ancestral polymorphism contributing to morph differentiation in different lake systems.

## Discussion

Here we have established a high-quality, scaffold-level genome assembly for the Arctic charr and provide gene annotation based on short and long-read RNA sequencing from multiple tissues including early developmental stages and cross-species comparisons. This provides a major advance for omic studies of this Arctic fish as well as for its utilization in aquaculture ^51,52^. We found that the nucleotide diversity within lakes was in the range 0.15-0.26%. Most of the sequence diversity we find among morphs within lakes is much older than the colonization and reflects founder effects and demographic history after colonization.

The population structure among Arctic charr from two Icelandic and two Norwegian lakes based on whole genome resequencing are largely consistent with previous results on Arctic charr from the Northern hemisphere based on more sparse sets of genetic markers (reviewed by Salisbury & Ruzzante, 2022). Firstly, populations clearly cluster by lakes rather than by morphs (Fig. 2). Our data are consistent with a scenario of early post-glacial isolation of these Arctic charr populations in the four lakes as impassable waterfalls formed downstream of the lakes ^19,35^. Secondly, three levels of genetic differentiation among sympatric morphs occur: (i) no genetic differentiation, (ii) genetic differentiation with incomplete reproductive isolation and (iii) strong genetic differentiation due to complete reproductive isolation. In Lake Vangsvatnet we find no genetic differentiation (Fig. 3d), implying that the phenotypic differentiation between the small benthic and planktivorous morphs in this lake is due to plasticity. In Lake Thingvallavatn (Fig. 3h) and Mývatn (Fig. 3f) we find clear genetic differentiation, localized in specific regions of the genome that well exceed the genomic background (Supplementary Fig. 2c, d-i), suggesting incomplete reproductive isolation and gene flow that tends to homogenize allele frequencies at neutral loci. Finally, in Lake Sirdalsvatnet we find strong genetic differentiation across the entire genome (Fig. 3a), consistent with complete or near complete reproductive isolation.

Why does the degree of genetic differentiation among morphs within lakes vary so much? Important factors include the time that has passed since a lake was colonized and chance events when critical steps towards differentiation occurred. Lake Vangsvatnet, estimated to be around 10,000 years old based on dating from a neighboring fjord ^53^, was likely colonized approximately 7,000 years BP (Jonsson 1982). In contrast, Lake Sirdalsvatnet is much older, likely been ice-free since 15,000 years BP. However, due to its location above high waterfalls, the exact timing of Arctic charr colonization remains uncertain. Another possibility is that genetically differentiated populations of charr colonized the same lake. However, the most important factor is likely ecological conditions within lakes ^54^. This is illustrated by the two Norwegian lakes showing drastically different population structures. In Vangsvatnet we found no genetic differentiation between morphs (Fig. 3c and d) whereas the corresponding morphs in Sirdalsvatnet showed strong, genome-wide divergence (Fig. 3a and b). The results are in line with differences in spawning habits (timing and area), where in Vangsvatnet (60 m deep) the two morphs partly co-occur in the spawning areas and may interbreed. Jonsson & Hindar (1982) reported that some small benthic females were found together with planktivorous males in shallow waters during spawning. In contrast, in Sirdalsvatnet (165 m deep) the majority of benthic and pelagic morphs spawn separately (the pelagic morph spawns at a depth of 0-32 m in November, whereas small benthic charr spawn at 55-70 m depth throughout the year ^55^, most of them in June-September, Supplementary Table 1) limiting possible gene flow. Morphs in both lakes exhibit habitat segregation for significant portions of the year. However, during periods of food surplus, this segregation seems to diminish in Vangsvatnet ^56^. Consistent with genetic differentiation, morphological studies suggest that the two morphs in Sirdalsvatnet differ more from each other in terms of the number of gill rakers, body size at sexual maturity, growth, and number of eggs per body size ^55^ than what the two morphs in Vangsvatnet do ^56^. In one of the deepest lakes in Norway, Tinnsjøen, in addition to dwarf benthic and planktivorous morphs, there is also a piscivorous morph, and a 5 cm long, colourless charr at the deepest part of the lake (460 m) ^22^, similar to the one found in the 288 m deep Gander Lake, Canada ^51,57^. This illustrates how variation in environmental conditions impacts sympatric differentiation in Arctic charr.

We noted clear genetic differentiation between morphs in Lake Mývatn (Iceland) implying that gene flow must be limited. Although the two morphs appear to overlap in timing and location of spawning (Supplementary Table 1), specifically within the main spawning ground of the generalist morph, microhabitat preferences for sites of spawning and size-assortative mate choice most likely limit inter-morph mating. Mývatn was formed around 2,300 years ago ^58^. Thus, the Arctic charr in Mývatn has a relatively recent colonization history compared to Thingvallavatn and the two Norwegian lakes. The Mývatn charr exhibits the highest nucleotide diversity among the populations under study (θ = 0.26%; Supplementary Table 7). The genome-wide changes observed in the Mývatn Arctic charr may include adaptive responses to various factors such as physical features of the benthic littoral zone and the extreme fluctuations of potential prey populations in the soft bottom and pelagic habitats of the main lake basins.

Thingvallavatn formed ∼10.000 years ago, first as a glacial lake and later, following large lava flows from the northern and eastern sides, as a spring fed lake ^59^. The lake was likely colonized by charr shortly thereafter. Our analysis of nucleotide diversity revealed reduced genetic diversity within the Thingvallavatn populations (θ = 0.15%) as compared to other lakes (θ = 0.24-0.26%). The most likely explanation for this reduced nucleotide diversity is a founder effect/bottleneck during colonization of the lake. Furthermore, geological events of the post glacial period, such as large lava flows, opening and collapsing of rifts connected to the lake and abrupt changes of water levels ^60^, could have impacted the genetic connectivity of morphs and lead to reductions in population size and genetic drift resulting in loss of genetic diversity. However, despite the lower nucleotide diversity, Thingvallavatn showed the most extensive phenotypic diversity with four distinct morphs present. This is most likely explained by the rich variety of habitats within this lake. The pattern of genetic differentiation among the four morphs present in Thingvallavatn seen in the genome-wide data (Fig. 3h) confirmed previous results based on much smaller sets of genetic markers ^7,15,35^. While genetic differentiation between the large benthic, small benthic and planktivorous morphs was clear, the genome-wide data showed that most piscivorous individuals were closely related to the planktivorous morph, but some showed intermediate or closer genetic affinities towards the large benthic morph (Fig. 3h), a pattern consistent with previous reports ^7,29,61,62^. A potential scenario is that the piscivorous charr in this lake have emerged and continue to emerge from the planktivorous fish via ontogenetic dietary shift to piscivory ^26^, a transition that leads to increased size at maturity thus opening the possibility of hybridization with the large benthic morph. However, we cannot exclude the possibility that large benthic males could be misclassified as piscivorous charr due to the similar head morphology (elongated lower jaw-hook). Some of these fish were found to be running in October when piscivorous charr were sampled ^7^.

A major advantage of the present study (whole genome sequencing and the use of a high-quality genome assembly) compared to previous studies using limited sets of markers, is that the genome-wide patterns of genetic differentiation become apparent. This resulted in the detection of as many as 12 putative inversions contributing significantly to the differentiation among morphs in Thingvallavatn (Figs 4 and 5). It is an open question whether these are true inversions or large haplotype blocks present due to reduced recombination and natural selection. It is challenging to identify inversion breakpoints based on short-read data since inversions are often flanked by repeats ^63^. Thus, further characterization of these putative inversions will require long-read sequencing of haplotypes. The Arctic charr is hereby added to a long list of fish species where putative or confirmed inversions play a significant role for intraspecies genetic differentiation. These include for instance Atlantic salmon where four confirmed inversions have been reported ^64–66^, Atlantic herring in which four confirmed inversions contribute to ecological adaptation related to water temperature ^67–69^, Atlantic silverside (13 putative inversions; ^70^ and European sprat, six putative inversions related to adaptation of this marine fish to brackish water bodies ^71^. Another example is the identification of large chromosomal inversions in rainbow trout associated with multiple adaptive traits ^72^ or geographic distribution ^73^. We propose that inversions may play a more prominent role for ecological adaptation in fish species than in birds and mammals, because fish are exposed to much more variable environmental conditions of particular importance during their early life stages when delicate developmental processes occur.

We noted a clear trend of differences in the level of nucleotide diversity between haplotypes at the 12 putative inversions detected in Thingvallavatn. The nucleotide diversity was consistently higher among the small benthic haplotypes compared with the large benthic haplotypes (Supplementary Table 3). Similarly, the pelagic haplotypes had consistently higher nucleotide diversity than the benthic haplotypes (Supplementary Table 6). In both cases, there was no significant difference between morphs in genome-wide nucleotide diversity, making it highly unlikely that differences in demographic history of the morphs explain the difference at the putative inversions. A reduced nucleotide diversity for one of the alleles at an inversion locus may reflect that this is the derived allele and/or that it has experienced a recent selective sweep.

An important question addressed in this study is the genetic basis for the characteristic sympatric differentiation in Arctic charr and to which extent genetic differentiation is based on ancestral polymorphisms shared among lake systems as well as genetic parallelism (i.e. how often the same or paralogous genes have contributed to adaptation). Firstly, it is clear that this species has an inherent ability to form phenotypically differentiated morphs even in the absence of genetic differentiation as noted for the Arctic charr in Vangsvatnet. This phenotypic plasticity is probably driven by the exploitation of different food resources (benthic versus pelagic in particular; ^74–76^. Secondly, genetic differentiation accumulates as some degree of reproductive isolation is established, e.g. based on spawning location and/or spawning time ^77^. Genetic adaptation is likely to be highly polygenic given the large number of genomic regions showing high differentiation in the current and previous studies ^3^, compared with the genomic background. This can be expected, given that Arctic charr morphs show phenotypic differentiation in multiple diverse traits such as feeding morphology, growth patterns, feeding behavior and spawning time ^18,78^. However, neoteny has been suggested as an important mechanism of morph formation as it influences many of the traits involving allometric growth and life history ^74,79^ and could in principle be driven by changes in one or few regulatory loci. Our comparison across lakes and morphs revealed that none of the 12 putative inversions showing strong genetic differentiation among the four morphs in Thingvallavatn segregated intact, with the same putative breakpoints, in any of the other three lakes. Further, the majority of the signals of selection outside the putative inversions do not appear to be shared among lakes. Thus, genetic differentiation of benthic and pelagic morphs in different lake systems show only limited genetic parallelism, for several possible reasons. The long time since the lakes were isolated, means independent evolution (no or limited gene flow between geographically isolated lakes), the lakes are quite different in topography and ecology and the phenotypes of the morphs differ considerably (lack of morphological parallelism). However, a notable exception is a 90 Kb region on scaffold 34, which represents a shared region of differentiation between benthic and pelagic morphs from Thingvallavatn and Mývatn, highlighting potential common adaptations across the Icelandic lakes (Supplementary Fig. 9). Additionally, we found clear indications of haplotype blocks of smaller segments within the putative inversion showing consistent morph associations among lakes from Iceland and Norway. One example is the putative inversion on scaffold 9 showing strong differentiation between small and large morphs in Thingvallavatn, where more than half of this genomic region replicates a similar pattern in Sirdalsvatnet (Supplementary Fig. 5). A possible scenario for the evolution of the putative inversions present in Lake Thingvallavatn is that they contain a combination of closely linked polymorphisms, some ancestral, that interact epistatically. This is a classical scenario for adaptive evolution of inversions ^80^. Haplotype sharing among lakes may in some cases represent ancestral inversions, but these inversion haplotypes have evolved by recombination which change the exact breakpoints. Inversions drastically reduce recombination but not completely. For instance, the inversion determining the satellite male morph in the ruff wader arose by a recombination event between an ancestral inversion and a wild-type chromosome ^81^. The results of the present study strongly suggest that whole genome resequencing of sympatric morphs from many lake systems in the Northern hemisphere presents a great opportunity to dissect the most important genetic factors that have contributed to the remarkable morph differentiation of Arctic charr across the Arctic region. Low-pass sequencing for determining population-specific allele frequencies combined with limited long-read sequencing, for the characterization of major structural variations, is a cost-effective approach to accomplish this.

Sympatric differentiation into two or more morphs after the colonization of lake systems with a paucity of fish biodiversity is a classical feature of salmonids including the Arctic charr ^3,82,83^. The whole genome duplication that occurred in an ancestor of salmonids about 100 Mya ^28^ may facilitate sympatric differentiation. There is evidence that the increased genome complexity in polyploids contributes to evolvability ^5^. Salmonids may tolerate loss of diversity due to founder effects when colonizing a new environment, because they have duplicated copies of many genes resulting in fixed heterozygosity when paralogs carry sequence differences, and new sequence variants can be created by genetic recombination between paralogs ^84^.

## Materials and methods

### Sampling for reference genome assembly and annotation

A large benthivorous charr from Lake Thingvallavatn was chosen for genome and transcriptome sequencing as part of the Pilot Project of the European Reference Genome Atlas (ERGA) initiative ^85^. LB charr were herded using a dragnet on the LB spawning grounds in Ólafsdráttur (DMM 64°13.8983’N, 21°03.1683’W) during the spawning season in late August. A single individual of the heterogametic male sex was caught by hand to serve as the ERGA reference sample. The fish was brought alive to the laboratory, euthanised with phenoxyethanol, photographed and immediately dissected for sampling from various tissues, strictly following ERGA guidelines. Tissue samples were snap-frozen in liquid nitrogen and stored at -80°C and shipped to the Earlham Institute and the University of Antwerp for processing. Voucher samples of fin and muscle were kept for archiving at the Icelandic Museum of Natural History. Sampling was done with license from the Icelandic Directorate of Fisheries and with permission from the Thingvellir National Park authority. Additional samples used for short-read RNA sequencing were from developmental time-series of offspring of LB charr parents from the same site, described in Matlosz *et al.* (2022), ENA project PRJEB45551).

### Reference genome assembly and annotation

#### Nucleic acid extraction

High-molecular weight DNA extraction was performed on snap-frozen spleen tissue using the Circulomics Nanobind Tissue Big DNA Kit. The extraction method was based on the Dounce protocol described in Circulomics protocol EXT-DHH-001. A total of 30 mg of tissue was used as input and split into five extractions with 6 mg input each to reduce viscosity. The eluted DNA was left at room temperature overnight, and periodically mixed 5 times with a wide-bore 200 μL tip the following day. RNA was extracted from seven tissue types from the same individual as used for the genome assembly. The Omega EZNA Total RNA Kit I (R6834-01) was used for the initial attempt from all tissues, and those that performed poorly (testis, muscle, and blood) were repeated using the EZNA Total RNA Kit II (R6934-01). For all solid tissues, 20-30 mg input was used per extraction. For blood, the entire frozen sample was thawed in an approximately equal volume of RNA-Solv buffer from the EZNA Total RNA Kit II. This mixture was then frozen at -80°C, and 250 μL was later taken for extraction. A 5-min GenoGrinder cycle with a 5mm steel bead was used for disruption of all tissue types, though the speed settings varied: 1000 rpm for gill, spleen, liver, and brain, and 1250 rpm for blood, testis, and muscle. To ensure small RNAs were not excluded from the final sample, the precipitation step following homogenization was performed with 100% ethanol instead of the recommended 70%.

#### Illumina RNA sequencing

Stranded mRNA-seq libraries were constructed at the Earlham Institute using the NEBNext Ultra II RNA Library prep for Illumina kit (NEB#E7760L), NEBNext Poly(A) mRNA Magnetic Isolation Module (NEB#7490) and NEBNext Multiplex Oligos for Illumina (96 Unique Dual Index Primer Pairs) (E6440S) and sequenced at a concentration of 10μM. 1 μg of RNA was purified to extract mRNA with a Poly(A) mRNA Magnetic Isolation Module. Isolated mRNA was then fragmented for 12 min at 94°C, and converted to cDNA. NEBNext Adaptors were ligated to end-repaired, dA-tailed DNA. The ligated products were subjected to a bead-based purification using Beckman Coulter AMPure XP beads (A63882) to remove most un-ligated adaptors. Adaptor Ligated DNA was then enriched by receiving 10 cycles of PCR (30 s at 98°C, 10 cycles of: 10 s at 98°C, 75 s at 65°C 5 min at 65°C, final hold at 4°C). The size of the resulting libraries was determined using Agilent High Sensitivity DNA Kit from Agilent Technologies (5067-4626) and the concentration measured with a High Sensitivity Qubit assay from ThermoFisher (Q32854). The final libraries were pooled equimolarly and quantified by qPCR. The pool was diluted down to 0.5 nM using EB (10mM Tris pH8.0) in a volume of 18μl before spiking in 1% Illumina phiX Control v3. This was denatured by adding 4μl 0.2N NaOH and incubating at room temperature for 8 min, after which it was neutralised by adding 5μl 400mM tris pH 8.0. A master mix of DPX1, DPX2, and DPX3 from Illumina’s Xp 2-lane kit was made and 63ul added to the denatured pool leaving 90μl at a concentration of 100pM. This was loaded onto a single lane of the NovaSeq SP flow cell (v1.5) using the NovaSeq Xp Flow Cell Dock before loading onto the NovaSeq 6000. The NovaSeq was run using NVCS v1.7.5 and RTA v3.4.4 and was set up to sequence 150bp PE reads. The data was demultiplexed and converted to fastq using bcl2fastq2.

#### Pacific Biosciences HiFi genome sequencing

Five spleen high-molecular weight extractions were combined to construct and sequenced two libraries at the Earlham Institute using the SMRTbell Express Template Prep Kit 2.0 (PacBio, P/N 100-983-900). In total, 23.4 µg was split into two aliquots and manually sheared with the Megaruptor 3 instrument (Diagenode, P/N B06010003) according to the Megaruptor 3 operations manual. Each aliquot underwent AMPure PB bead (PacBio, P/N 100-265-900) purification and concentration before undergoing library preparation using the SMRTbell Express Template Prep Kit 2.0 (PacBio, P/N 100-983-900). The HiFi libraries were prepared according to the HiFi protocol version 03 (PacBio, P/N 101-853-100) and the final libraries were size fractionated using the SageELF system (Sage Science, P/N ELF0001), 0.75% cassette (Sage Science, P/N ELD7510). The libraries were quantified by fluorescence (Invitrogen Qubit 3.0, P/N Q33216) and the size of the library fractions were estimated from a smear analysis performed on the FEMTO Pulse System (Agilent, P/N M5330AA). The libraries were sequenced on the Sequel IIe across eight Sequel II SMRT cells 8M. The parameters for sequencing per SMRT cell were: Adaptive loading default settings, 30-h movie, 2-h pre-extension time, 80-90pM on plate loading concentration. The loading calculations for sequencing were completed using the PacBio SMRTLink Binding Calculator 10.2. Sequencing primer v5 was annealed to the adapter sequence of the HiFi libraries. The libraries were bound to the sequencing polymerase with the Sequel II Binding Kit v2.2 (PacBio, P/N 102-089-000). Calculations for primer and polymerase binding ratios were kept at default values for the library type. Sequel II DNA internal control 1.0 was spiked into each library at the standard concentration prior to sequencing. The sequencing chemistry used was Sequel II Sequencing Plate 2.0 (PacBio, P/N 101-820-200) and the Instrument Control Software v 10.1.0.125432.

#### Pacific Biosciences Iso-Seq sequencing

The libraries were constructed and sequenced at the Earlham Institute starting from 231-348ng of total RNA per sample. Reverse transcription cDNA synthesis was performed using NEBNext Single Cell/Low Input cDNA Synthesis & Amplification Module (NEB, E6421) Each cDNA sample was amplified with barcoded primers for a total of 12 cycles. The barcoded cDNA samples were pooled equimolar into a 3-plex and 4-plex before SMRTbell library construction. Each library pool was prepared according to the guidelines laid out in the Iso-Seq protocol version 02 (PacBio, 101-763-800), using SMRTbell express template prep kit 2.0 (PacBio, 102-088-900). The library pool was quantified using a Qubit Fluorometer 3.0 (Invitrogen) and sized using the Bioanalyzer HS DNA chip (Agilent Technologies, Inc.). Each Iso-Seq pool was sequenced on the Sequel IIe instrument with one Sequel II SMRT Cell 8M per pool. The parameters for sequencing were diffusion loading, 30-h movie, 2-h immobilisation time, 2-h pre-extension time, 60pM on plate loading concentration. The loading calculations for the Iso-Seq library pool using the PacBio SMRTlink Binding Calculator v 11.1.0.154383 and prepared for sequencing applicable to the library type. Sequencing primer v4 was annealed to the Iso-Seq library pool and complexed to the sequencing polymerase with the Sequel II binding kit V2.1 (PacBio, 101-843-000). Calculations for primer to template and polymerase to template binding ratios were kept at default values for the library type. Sequencing internal control complex 1.0 (PacBio, 101-717-600) was spiked into the final complex preparation at a standard concentration before sequencing for all preparations. The sequencing chemistry used was Sequel II Sequencing Plate 2.0 (PacBio, 101-820-200) and the Instrument Control Software v11.0.1.162970.

#### Dovetail Omni-C sequencing

A Hi-C library was prepared using Dovetail Genomics’ Omni-C kit, following the manufacturer’s protocol (v1.0) at the University of Antwerp. A liver tissue sample was manually homogenized with a micro pestle and needle and syringe before proceeding with the Hi-C library preparation steps. Library fragment size distribution and concentration were assessed using High Sensitivity D5000 ScreenTapes with a TapeStation 4150 (Agilent Technologies) and a Qubit Fluorometer (Thermo Fisher Scientific, Waltham, MA, USA), respectively. The Omni-C library was denatured and loaded for paired-end sequencing on an Illumina NovaSeq 6000 system using the S1 Reagent Kit v1.5 (300 cycles) at the University of Florence. We set a single index running mode to 6:151:151:0 bp cycles. Demultiplexing and conversion of sequencing data from bcl to fastq format were performed using bcl2fastq version 2.20 (Illumina).

#### Ploidy and genome size estimation

GenomeScope 2.0 ^87^ was used to generate a reference-free k-mer spectra profile and a smudgeplot to predict the ploidy of the species. First a meryl database was generated using a k-mer size of 21 (-k 21). The smudge plot was generated using kmers with a count > 200 and <3200.

#### Contig assembly

Hifiasm v0.18.5 ^88,89^ was used to assemble the PacBio HiFi reads into a contiguous draft genome, using Omni-C reads to phase the genome into two phased genomes. A contamination screening was realized to the contigs of both assemblies using diamond v2.0.9 ^90^ blastx against the NCBI NR database (downloaded 29/03/2023) with the options --fast and --strand both to map against the positive and negative strands. The resulting hits were used as input for blobtools v1.0.1 ^91^ to generate a blobplot and a results table with the most likely phylogeny of each contig. Most of them were classified as Actinopterygii as expected for this species with some hits to other phylum. These contigs were manually blasted and the top hits were always from the Salmonidae family. Mito-Hi-Fi v2.2 ^92^was used to identify, extract and annotate the mitochondria from both assemblies. Only the mitochondria from the primary haplotype were kept for further analysis.

#### Scaffolding

YAhS v1.2 ^93^ was used to scaffold the contigs using the Omni-C reads using a resolution of 1kb.

#### Manual curation

The final screened assembly was curated by the Genome Reference Informatics at the Wellcome Sanger Institute using the Omni-C data to generate a contact map and break/join scaffolds that had enough information to be associated as one DNA molecule.

#### Assembly k-mer spectra analysis

The quality and completeness of the assembly was assessed using merqury v1.3 ^94^ using the meryl database created for the ploidy and genome size estimation step.

#### Repeat masking

Repeats were identified and masked using RepeatModeler v1.0.11 ^95^ and RepeatMasker v4.0.72 ^96^ via eirepeat v1.3.4 (https://github.com/EI-CoreBioinformatics/eirepeat).

#### Gene prediction

Gene models were annotated via the Robust and Extendable eukaryotic Annotation Toolkit (REAT, https://github.com/EI-CoreBioinformatics/reat) and Minos (https://github.com/EI-CoreBioinformatics/minos). The REAT workflow consists of three submodules: transcriptome, homology, and prediction. The transcriptome module utilised Illumina RNA-Seq data, mapping reads to the genome with HISAT2 v2.1.0 ^97^ followed by the identification of high-confidence splice junctions identified with Portcullis ^98^. The aligned reads were assembled for each tissue with StringTie2 v1.3.3 ^99^ (Kovaka et al., 2019) and Scallop v0.10.2 ^100^. From the combined set of RNA-Seq assemblies a filtered set of non-redundant gene-models were derived using Mikado https://github.com/EI-CoreBioinformatics/mikado ^101^. The REAT homology workflow was used to generate gene models based on alignment of protein sequences using Spaln2 ^102^ and miniport ^103^ from 10 related species previously annotated (Supporting Information Table SX) and a set of proteins downloaded from UniProt including all the proteins from the Actinopterygii class (taxid:7898). The prediction module generated evidence guided models based on transcriptome and protein alignments using AUGUSTUS v3.4.0 ^104^ (Stanke & Morgenstern, 2005) with three alternative configurations and weightings of evidence (see configs), Helixer ^105^ and EvidenceModeler v1.1.1 ^106^. Gene models from annotations of closely related organisms: *Salvelinus* spp., *Salvelinus namaycush*, *Salvelinus fontinalis* (GCF_002910315.2, GCF_016432855.1, GCF_029448725.1) were projected via Liftoff v1.5.1 ^107^, and filtered via the multicompare script from ei-liftover pipeline (https://github.com/lucventurini/ei-liftover) ensuring only models with consistent gene structures between the original and transferred models were retained. The filtered Liftoff, REAT transcriptome, homology and prediction gene models were used in MINOS v1.9 (https://github.com/EI-CoreBioinformatics/minos) to generate a consolidated gene set with models selected based on evidence support and their intrinsic features (see config directory). Confidence and biotype classification was determined for all gene models based on available evidence, such as homology support and expression (as defined in config_file). Transposable element gene classification was based on overlap with identified repeats (>40 bp repeat overlap).

### Samples for whole genome resequencing

Arctic charr samples for whole genome analyses were obtained from two lakes in Iceland: Thingvallavatn and Mývatn and from two lakes in Norway: Vangsvatnet and Sirdalsvatnet (Fig. 1). Samples from the two Norwegian lakes were from a previously published study ^19^.

The four charr morphs of Lake Thingvallavatn were sampled at the respective spawning grounds using gillnets of various mesh sizes. LB charr were sampled at Ólafsdráttur. PL, Pi and SB charr were sampled on both sides of the Mjóanes peninsula. The sampling effort and DNA extraction is described in detail in Xiao et al. (2024).

Samples of the large generalist morph (LG) from Lake Mývatn were caught in gillnets in the main basin of the lake as part of regular fisheries surveys by the Icelandic Marine and Freshwater Research Institute and kindly provided by Guðni Guðbergsson. Sampling for the SB morph (locally known as “Krús”) was conducted at Kálfaströnd, in an area of cold-water springs (N 65°33.841 W 16°56.681). The fish were caught by electrofishing, anesthetized using phenoxyethanol at low concentration and measured, weighed and photographed. Small clippings from the caudal fin were stored in ethanol and the fish were then released. Many fish from Kálfaströnd were 0+ and 1+ juveniles that could not be assigned to morph based on morphology. As the cold spring area is a known habitat harbouring the SB-morph all the fish caught at Kálfaströnd were pre-assigned to that morph. Morph assignment was later revised based on the photographs and genetic data (Supplementary Fig 2).

Detailed information on the number of samples from each morph at each sample location are presented in Supplementary Table 1.

### Whole genome resequencing

Extracted genomic DNA was available for the Icelandic samples ^29^. Briefly, ethanol-preserved tissue from fin clips or muscle was rehydrated and digested with proteinase K followed by DNA isolation using standard phenol/chloroform extraction. Muscle tissue preserved in 99% ethanol was used for DNA extraction from the Norwegian samples. Genomic DNA was extracted using the Monarch Genomic DNA Purification Kit (New England Biolabs). The purity of the DNA was assessed with Nanodrop 1000, and the DNA quantity was assessed using the Qubit dsDNA BR Assay Kit (Thermo Fisher Scientific) and a Tecan microplate reader. The DNA was diluted to a final concentration of 10 ng/μL for library preparation. Custom Tn5-based libraries ^108^ with a target size of 350 bp were constructed for each DNA sample (n = 283). All libraries were individually barcoded and pooled. The TapeStation D1000 (Agilent Technologies) was used to visualize library size. The library pool was sequenced on two Illumina NovaSeq S4 lanes.

The Illumina raw reads were mapped to the Arctic charr assembly (ENA Project PRJEB76174), developed as part of this study. Mapping was done using “bwa mem -M” v0.7.17 (Li, 2013). The resulting alignments were sorted using Samtools v1.10 (http://www.htslib.org/) and finally processed with MarkDuplicates from PicardTools v1.92 (https://broadinstitute.github.io/picard/). The average sequencing depth per individual was 2.1±0.9 (range: 0.22-6.11).

### Genotype likelihood estimation and quality control

Due to the limitations of the low sequencing coverage approach for genotype calling, we estimated genotype likelihoods using ANGSD v0.933 ^109^. The genotype likelihoods were used to estimate allele frequencies for each morph (n = 22-40, Supplementary Table 1) or population (n = 52-111, Supplementary Table 1). Depending on the analysis, the following parameters were used: “uniqueOnly 1 -remove_bads 1 - only_proper_pairs 0 -trim 0 -GL 2”. We used “– doMajorMinor 4 –minMaf 0.05” options to generate the list of positions with minor allele frequency (MAF) > 5%. To generate the high-confidence variants for the analysis we supplied “-SNP_pval 1e-6” to ANGSD. The total set of variant positions across the entire data set was ∼12 million, of which 1.5 million out of each were on unplaced scaffolds. The list of polymorphic positions was supplied as “-sites” argument for genotype likelihood, population-or morph-wise allele frequencies estimation and association study. Principal component analysis (PCA) and calculations of ancestry proportions and pairwise Nei distances were performed based on a reduced set of SNPs. This dataset was obtained by selecting one SNP for every 10 in the original dataset and applying a MAF cutoff of 5%, resulting in varying numbers of retained SNPs: ∼1 million SNPs for all lakes combined, 0.73 million SNPs for Sirdalsvatnet, 0.65 million SNPs for Vangsvatnet, 0.51 million SNPs for Mývatn, and 0.58 million SNPs for Thingvallavatn. Nucleotide diversity calculation used all observed positions. All data were obtained for 40 scaffolds, while unplaced scaffolds were summarized into a NA scaffold.

### Analysis of population structure

To examine population structure and genetic divergence, we ran PCA using PCAngsd v1.11 ^110^ for the data sets across and within lakes. The built-in R function “eigen()” was used to extract eigenvalues and eigenvectors, “ggplot2” package was used to plot the data.

We calculated individual admixture proportions using PCAngsd v1.11 ^110^ and NGSadmix v. 32 ^111^ modules from ANGSD. PCAngsd was used for all lakes combined and determines the optimal number of clusters (K) empirically based on the major principal component (PC) loadings. In contrast, NGSadmix was applied within each lake separately, with a predefined K. To identify the best-supported number of clusters (K), we evaluated log-likelihoods and Frobenius errors across a range of K values (1 to 10) following Meisner & Albrechtsen (2018). The optimal K was determined as the value that maximized the log-likelihood and minimized the Frobenius error.

A neighbor-joining (NJ) genetic distance tree of samples from a covariance matrix was derived based on individual allele frequencies using the “-tree” flag in PCAngsd ^110^. The NJ tree visualization was done using an online tool iTOL (https://itol.embl.de/, Ciccarelli et al., 2006).

### Genetic diversity

To characterize nucleotide diversity for each sampling location and morph, we calculated the average number of pairwise differences between sequences (θ). All θ were calculated based on allele frequencies generated without MAF cutoffs because MAF cutoffs introduce bias in population-based frequency diversity estimates. The following command implemented in ANGSD v.0.933 software ^109^ was used to calculate allele frequencies per sample group originating from the same lake or separately from each morph within a lake:

“angsd -bam $POPULATION -anc $REFGENOME -fai $REF_INDEXED -doSaf 1 -GL 2 -P 10 -out output -doCounts 1 -setMinDepth 15 -setMaxDepth 1000 -setMinDepthInd 0.25 - minMapQ 30 -minQ 20 -remove_bads 1 -minInd 10 -uniqueOnly 1 -dumpCounts 2”.

The filters above select sites with high-quality sequencing depth and alignment, and to ensure the selected sites have sequence information in at least 10 individuals per population (*-minInd 10*). After that we used the ANGSD realSFS command to calculate a 2-dimensional folded site frequency spectrum (2D-SFS). This was used as a prior to calculate diversity statistics with the saf2theta command. Pairwise nucleotide diversity was calculated in 5 kb non overlapping windows. Obtained values were averaged per population and divided by the number of sites per window to recover an unbiased estimate of nucleotide diversity as described ^113^.

Finally, population differences were assessed using pairwise Fst. The resulting 2D-SFS from the previous step were used as a prior to estimate Fst statistics with the ANGSD realSFS fst index command. To perform a sliding window analysis, Fst values were computed with a window size of 20 kb and a step size of 10 kb using the realSFS fst stats2 command, with the options -win 20000 and -step 10000. Considering that estimation of Fst relies on the presence of enough genetic variation (polymorphic sites) to make a meaningful comparison between populations, the first five largest scaffolds were used in the Fst scan (∼113-165 million sites). In the case of pairwise Fst estimates between morphs within each lake, scaffolds without putative inversions were used (e.g, scaffolds 2, 6, 7, 10 and 11 with ∼21-152 million sites). The obtained Fst values for each population or morph pair were averaged across the scaffolds.

### Genome wide screen for genetic differentiation

To identify genomic regions showing genetic differentiation between populations we used a genome-wide association study (GWAS). GWAS was performed in ANGSD using the “-doAsso’’ flag along with “-Pvalue 1” to obtain *P*-values in the final output file. This association study is based on individual genotype likelihoods and takes all uncertainty of the generated data into account ^109^.The method used compares allele frequencies between all pairs of morphs within a lake. The genome-wide significance threshold was calculated using a Bonferroni correction with a significance level at *α =* 10^−3^ and *α =* 10^−8^ using the following formula: Bonferroni = -log10(*α*/N), where N is the number of SNPs.

### Analysis of putative structural variants

SNPs showing strong genetic differentiation exceeding the Bonferroni-corrected significance threshold (*α =* 10^−8^) were selected as diagnostic markers. These markers were used to investigate haplotype sharing across morphs within and across lakes. A haplotype was defined as a set of variants closely located (up to 50 kb) along a single chromosome, with strong linkage disequilibrium (LD) estimated based on genotype likelihoods. LD was quantified using ngsLD/1.1.1, with an *R*^2^ threshold of 0.5 to identify linked variants. The 50 kb window size was selected based on empirical evidence, reflecting the expected genomic linkage within the studied populations. Nucleotide diversity (θ) of each haplotype for the putative inversions was estimated using ANGSD as described above. In addition, based on calculating the posterior probability of the genotypes we run genotype calling using “-doGeno” and “-doPost” flags in ANGSD. We summed the called genotypes supporting the reference and the alternative alleles for each sample, selected the most common allele in a reference group, and kept genotypes for only those positions. Based on the genotype distribution we formed three groups (homozygous major, homozygous minor and heterozygous) and compared θ distributions amongst them using Wilcoxon rank test for a difference in means in R (v4.3.2) ^114^.

### Analysis of haplotype sharing outside putative structural variants

To analyze haplotype sharing (see the description above) across morphs from different lakes outside the putative structural variants, we compared allele frequency and predicted genotype distributions at diagnostic markers as described above. To validate the presence of shared haplotype, individuals clustered as homozygous major and homozygous minor were then used for the PCA analysis and NJ genetic distances tree using PCAngsd ^110^.

### Functional annotation of gene models

The gene models in the reference genome (described above) initially only included gene IDs and lacked gene symbols and descriptions. To address this and enhance interpretations of the results, we utilized the functional gene annotation pipeline provided by the National Bioinformatics Infrastructure Sweden (NBIS, Binzer-Panchal et al., 2021) to obtain the missing information. Briefly, the nucleotide sequence of each annotated gene from the genome sequence in FASTA format using the gene coordinates from the GFF annotation file. The nucleotide sequence per gene was then translated into amino acid sequence using Another Gtf/Gff Analysis Toolkit (AGAT v.1.3.2) ^116^. The amino acid sequence was compared against a reference protein database to infer which protein corresponds to each gene using BLASTp v.2.15.0+ ^117^ with blast e-value = 1e-6. As a reference we used the reviewed Vertebrates protein database available from UniProtKB/SwissProt in July 2021 (version 2021_03). Gene function annotation was conducted using STRING v12 (https://string-db.org/), with Homo sapiens selected as the reference gene database for the analysis. Finally, we annotated the nearest gene (within ±5 kb) and assessed the variant effects (such as missense, synonymous, upstream, downstream, or intergenic) for the of the most differentiated SNPs (*P* <1×10^−15^) identified in each putative inversion or shared region of divergences using SnpEff v.4.1 ^118^.

## Acknowledgments

Short-read Illumina sequencing was performed by the SNP&SEQ Technology Platform in Uppsala. The facility is part of the National Genomics Infrastructure (NGI) Sweden and Science for Life Laboratory. The SNP&SEQ Platform is also supported by the Swedish Research Council and the Knut and Alice Wallenberg Foundation. Computational infrastructure was provided by the National Academic Infrastructure for Supercomputing in Sweden, partially funded by the Swedish Research Council through grant agreement no. 2022-06725. The frozen tissue bank storing of the Norwegian Arctic charr samples used here is maintained with support from the Department of Zoology, Stockholm University, the Swedish Research Council, and Formas, the Swedish Agency for Marine and Water Management, the Swedish Environmental Protection Agency. The authors acknowledge the work of the ERGA Pilot Project coordinators, Ann Mc Cartney, Giulio Formenti and Alice Mouton, in making this work possible by establishing the necessary sample metadata collection infrastructure, including the early establishment of sample manifest collection process, the ERGA manifest Github repository, the in-kind reagents from commercial companies, as well as with their constant coordination effort as part of the ERGA Pilot Project daily activities. We thank Alex Durant and Jonathan MD Wood for their contribution to the development of the genome assembly.

## Author contributions

KG and ZOJ led the development of the reference assembly. LA led whole genome resequencing. KK was responsible for the population genomics analysis. AC constructed the Tn5 libraries for whole genome sequencing. MP contributed to the population genomics analysis. ZOJ, SS and AP sampled the Icelandic lakes with the exception of the large morph of Mývatn. ZOJ, SS and AP dissected the LB voucher sample for transcriptome sequencing. HX performed DNA extractions from all Icelandic samples used for low pass sequencing. GD and HL prepared the Hi-C libraries under the supervision of HS. KK and LA wrote the paper with input from other coauthors. All authors approved the manuscript before submission.

## Data and materials availability

All custom code used for processing and analysis of the whole-genome re-sequencing data is available in https://github.com/LeifAnderssonLab/Arctic_charr_morphs. Whole genome re-sequencing data in this study have been deposited in the Sequence Read Archive under BioProject accession number PRJNA1175689. Raw genomic reads used for reference genome assembly and annotation, assembly sequences and reference annotations are available from the European Nucleotide Archive under Project PRJEB76174.

