## Supplementary materials for "Whole genome sequencing reveals how plasticity and genetic differentiation underlie sympatric morphs of Arctic charr"

**List of Supplementary Materials**

**Supplementary Fig. 1. Syntenic regions identified between assembly scaffolds and the Canadian Arctic Char high-density linkage map.** GBS sequences assigned to sex-specific linkage groups described in [1] were mapped to the 40 chromosome-level scaffolds in the assembly. Linkage groups for the male (A) and female (B) maps are shown in rows while scaffolds are shown in columns. The figures in each cell represent the total number of GBS sequences shared between the corresponding linkage group and scaffolds. Syntenic blocks supported by three or more GBS sequences are highlighted in black. Empty cells indicate a lack of shared sequences.

**Supplementary Fig. 2 Genetic differentiation among Arctic charr morphs in Lake Mývatn**. **(a)** Scores of individuals along PC1 and PC2, and **(b)** ancestry proportions of individuals assuming two clusters (K=2), estimated based on a downsampled set of 0.51 million SNPs (MAF > 0.05). The pre-assignment of morphs was based on sampling habitats. Morph abbreviations: Large generalist (LG) and Small benthic (Krús, SB).

**Supplementary Fig. 3 Genomic differentiation between morphs of Arctic charr from four lakes.** Genome scan based on estimated allele frequencies for individual SNPs. (**a)** Sirdalsvatnet, (**b)** Vangsvatnet, (**c**) Mývatn, and (**d–i**) Thingvallavatn. The x-axis represents the scaffolds, and the y-axis represents the significance value (-log10(*P*)-value) per SNP. Each dot corresponds to a single SNP. The horizontal red line indicates the significance threshold based on Bonferroni correction with the adjusted significance level for $\alpha = 10^{-3}$. Morph abbreviations for Sirdalsvatnet and Vangsvatnet: Dwarf benthic (DB) and Large pelagic (LP); for Mývatn, Large generalist (LG) and Small benthic (Krús, SB), and Thingvallavatn, Piscivorous (Pi), Planktivorous (PL), Large benthivorous (LB) and Small benthivorous (SB).

**Supplementary Fig. 4 Admixture analysis of individuals from Lake Thingvallavatn. (a)** Ancestry proportions generated in NGSAdmix based on a downsampled set of 0.58 million SNPs (MAF > 0.05). Morph abbreviations: Piscivorous (Pi), Planktivorous (PL), Large benthivorous (LB), and Small benthivorous (SB). (**b)** Log-likelihoods for 1–10 clusters in the admixture analysis. K = 3 was determined to best represent the genetic structure in the lake, aligning with the three distinct groups observed in the PCA plot (Fig. 3g**)**. Increasing K beyond 3 resulted in only minor changes in admixture proportions, suggesting that three main genetic groups capture the primary population structure within this lake.

**Supplementary Fig. 5 Predicted genotypes based on genotype likelihoods from diagnostic markers at four putative inversions distinguishing large and small benthic morphs in Thingvallavatn.** Genotype distributions sorted by lake and morph are shown for the putative inversion regions on scaffolds 4, 5, 9, and 17. Morph abbreviations for Sirdalsvatnet and Vangsvatnet: Dwarf benthic (DB) and Large pelagic (LP); for Mývatn, Large generalist (LG) and Small benthic (Krús, SB), and Thingvallavatn, Piscivorous (Pi),

Planktivorous (PL), Large benthivorous (LB) and Small benthivorous (SB). Individuals are colored according to their estimated genotype. The black brackets indicate a shared pattern of differentiation between morphs.

**Supplementary Fig. 6 Predicted genotypes based on genotype likelihoods from diagnostic markers at six putative inversions distinguishing Arctic charr benthic (large and small benthivorous) and pelagic (piscivorous and planktivorous) morphs in Thingvallavatn.** Genotype distributions sorted by lake and morph are shown for the putative inversion regions on scaffolds 1, 3, 8, 9, 14, and 40. Morph abbreviations for Sirdalsvatnet and Vangsvatnet: Dwarf benthic (DB) and Large pelagic (LP); for Mývatn, Large generalist (LG) and Small benthic (Krús, SB), and Thingvallavatn, Piscivorous (Pi), Planktivorous (PL), Large benthivorous (LB) and Small benthivorous (SB). Individuals are colored according to their estimated genotype. The black brackets indicate a shared pattern of differentiation between morphs.

**Supplementary Fig. 7 Allele frequency of diagnostic SNPs outside the putative inversions selected based on the small/large benthivorous contrast of Arctic charr morphs from Thingvallavatn.** Each column title denotes a scaffold, with each value representing a diagnostic marker. The colors of the column annotation panel denote haplotype. Haplotypes were defined based on a windowed approach, where the maximum distance allowed between adjacent single nucleotide polymorphisms (SNPs) within the same haplotype was set to 50 kilobases (kb). This threshold was chosen to capture localized linkage disequilibrium patterns, ensuring that SNPs within each haplotype block were tightly linked and likely to be inherited together. Only regions containing more than four SNPs were included to ensure sufficient density for linkage analysis. Each of the presented regions ranges in size from 0.116 to 0.146 Mb. SNPs were tracked based on the most common allele in the Small benthivorous morph from Thingvallavatn. Allele frequencies (AF) of these alleles are indicated by the color code, grey color indicates missing data. Morph abbreviations for Sirdalsvatnet and Vangsvatnet: Dwarf benthic (DB) and Large pelagic (LP); for Mývatn, Large generalist (LG) and Small benthic (Krús, SB), and Thingvallavatn, Piscivorous (Pi), Planktivorous (PL), Large benthivorous (LB) and Small benthivorous (SB).

**Supplementary Fig. 8 Allele frequency of diagnostic SNPs outside the putative inversions selected based on the benthic/pelagic contrast of Arctic charr morphs from Lake Thingvallavatn.** Each row title denotes a scaffold, with each value representing a diagnostic marker. The colors on the left annotation panel denote haplotype. Haplotypes were defined as described in Supplementary Fig. 6. Each of the presented regions ranges in size from 0.167 to 0.314 Mb. SNPs were tracked based on the most common allele in the benthic morph from Thingvallavatn. Allele frequencies (AF) of these alleles are indicated by the color code, grey color indicates missing data. Morph abbreviations as in Supplementary Fig. 6.

**Supplementary Fig. 9 Genetic differentiation at a region on scaffold 34 among Arctic charr morphs in Mývatn and Thingvallavatn. (a)** Predicted genotypes based on genotype likelihoods from diagnostic markers at this region sorted by lake and morph.

Morph abbreviations as in Supplementary Fig. 6. Individuals are colored according to their estimated genotype. Each value in the column represents a diagnostic marker. SNPs were tracked based on the most common allele in benthic morphs from Thingvallavatn. **(b)** Zoom-in profile of the genome-wide scan based on estimated allele frequencies for individual SNPs and linkage disequilibrium represented as R2 among genotypes in the diagnostic region on scaffold 34 (18.36-18.45 Mb). **(c)** PCA plot showing individual clustering of samples homozygous for benthic or pelagic haplotype from Mývatn and Thingvallavatn within the diagnostic region (scaffold 34: 18.36-18.45 Mb). **(d)** Neighbor-joining tree of samples homozygous for benthic or pelagic haplotype from Mývatn and Thingvallavatn lakes within the diagnostic region (scaffold 34: 18.36-18.45 Mb) and **(e)** outside the diagnostic region (scaffold 34: 0.01-18.3 Mb and 18.5 - 40.7 Mb).

**Supplementary Table 1** Sample metadata, geological context, and history of the studied lakes where distinct Arctic charr morphs occur, including spawning times of the study populations. The Norwegian samples were collected from 1980 to 1984 and were the same as those used by Hindar et al. (1986) and they are maintained in a frozen tissue bank kept by L.L and N.R. at the Department of Zoology, Stockholm University. The samples from Thingvallavatn were collected in 2016-2018. The samples of LG-charr from Mývatn were taken in 2014 from a fisheries survey by the Freshwater Research Institute, and the samples of SB (Krús) charr were collected in 2015.

**Supplementary Table 2** Number of scaffolds and SNPs where at least one marker exceeds the Bonferroni corrected significance level ($\alpha = 10^{-3}$) identified per each contrast between Arctic charr morphs from the four lakes. Morph abbreviations for Sirdalsvatnet and Vangsvatnet: Dwarf benthic (DB) and Large pelagic (LP); for Mývatn, Large generalist (LG) and Small benthic (Krús, SB), and Thingvallavatn, Piscivorous (Pi), Planktivorous (PL), Large benthivorous (LB) and Small benthivorous (SB).

**Supplementary Table 3** Genotype distribution at four putative inversions showing genetic differentiation between the small and large benthivorous and large benthic morphs present in Thingvallavatn.

**Supplementary Table 4** Nucleotide diversity (θ) assessed across the whole genome and at putative inversion regions among Arctic charr morphs from Thingvallavatn homozygous for small and large benthivorous haplotype

**Supplementary Table 5** Genotype distribution at eight putative inversions showing genetic differentiation between the benthic (B) and pelagic (P) morphs present in Thingvallavatn.

**Supplementary Table 6** Nucleotide diversity (θ) assessed across the whole genome and at putative inversion regions among Arctic charr morphs from Lake Thingvallavatn homozygous for benthic or pelagic haplotype.

**Supplementary Table 7** Nucleotide diversity (θ) assessed across the whole genome among Arctic charr morphs and populations. Morph abbreviations for Sirdalsvatnet and Vangsvatnet: Dwaf benthic (DB) and Large pelagic (LP); for Mývatn, Large generalist (LG) and Small benthic (Krús, SB), and Thingvallavatn, Piscivorous (Pi), Planktivorous (PL), Large benthivorous (LB) and Small benthivorous (SB).

**Supplementary Table 8** Genotype distribution at locus 18.36-18.45 Mb of scaffold 34 showing genetic differentiation between morphs homozygous for benthic and pelagic haplotype present in Thingvallavatn and Mývatn.

**Supplementary Table 9** Gene list for regions within 5 kb upstream and 5 kb downstream of putative inversions distinguishing small vs large benthic morphs in Thingvallavatn. (added in a separate excel file Supplementary tables 9-12)

**Supplementary Table 10** Gene list for regions within 5 kb upstream and 5 kb downstream of putative inversions distinguishing benthic and pelagic morphs from Thingvallavatn. (added in a separate excel file Supplementary tables 9-12)

**Supplementary Table 11** Gene list for regions within 5 kb upstream and 5 kb downstream of Scaffold 34 (18.36–18.45 Mb), associated with haplotypes distinguishing benthic and pelagic morphs in Mývatn and Thingvallavatn. (added in a separate excel file Supplementary tables 9-12)

**Supplementary Table 12** Summary of the most differentiated SNPs ($P < 1\times10^{-15}$) for each diagnostic region, including their nearest gene, and relative position to genes (e.g., missense, synonymous, upstream, downstream, or intergenic) as determined by snpEff. (added in a separate excel file Supplementary tables 9-12)

**Supplementary Fig. 1. Syntenic regions identified between assembly scaffolds and the Canadian Arctic Char high-density linkage map.** GBS sequences assigned to sex-specific linkage groups described in [1] were mapped to the 40 chromosome-level scaffolds in the assembly. Linkage groups for the male (A) and female (B) maps are shown in rows while scaffolds are shown in columns. The figures in each cell represent the total number of GBS sequences shared between the corresponding linkage group and scaffolds. Syntenic blocks supported by three or more GBS sequences are highlighted in black. Empty cells indicate a lack of shared sequences.

**A**

| | 1 | 2 | 3 | 4 | 5 | 6 | 7 | 8 | 9 | 10 | 11 | 12 | 13 | 14 | 15 | 16 | 17 | 18 | 19 | 20 | 21 | 22 | 23 | 24 | 25 | 26 | 27 | 28 | 29 | 30 | 31 | 32 | 33 | 34 | 35 | 36 | 37 | 38 | 39 | 40 |
|---|---|---|---|---|---|---|---|---|---|---|---|---|---|---|---|---|---|---|---|---|---|---|---|---|---|---|---|---|---|---|---|---|---|---|---|---|---|---|---|---|
| AC-1+21m | | | | 1 | 1 | | | | | | 93 | | | 1 | | 1 | | | | | 1 | | | 1 | 42 | | | | | | 1 | | | | | | | 45 | | 1 |
| AC-2m | | | | | | | | | | | | | | | 1 | | | | | 1 | 93 | | | | | | | | | | | | 1 | | | | | | | |
| AC-3m | 1 | | | 2 | | | | | 68 | | | | | | | | | | | | | | | | | | | | | | | | | | | | | | 3 | |
| AC-4p-m | | | | | | | 1 | 1 | | | | | | | | | | 1 | | | | | 3 | | | | | | 1 | 39 | | | | | | | | | | |
| AC-4q-m | 89 | | 1 | | | | | | | | | | | | | | | | | | 1 | | | | | | | | | | | | | 1 | | | | | | |
| AC-5m | | | | | | 1 | | | 1 | | 1 | | | 1 | | | | | | | | 88 | | | | | | | | | | | | | | | | | | |
| AC-6m | | | | 1 | 90 | 1 | | | | | 1 | 1 | | | | | | | 1 | | | | | | | | | | | | | | | | 1 | | | | | |
| AC-7m | | | | | | | | | | | | | | | | | | | | | | | | | | | | | 46 | | | | | | | | | | | |
| AC-8m | | | | | | 58 | | | | | | | | | | | | | | | | | | | | | | | | | | | | | | | | | | |
| AC-9m | 2 | | | 2 | | | | 1 | | | | | | | | | | | | | | | | | | | 72 | | | | | | | | | | | | | |
| AC-10m | | 1 | | 6 | | | 1 | | | | | | | | | | | | | | | | | | | | | 1 | | | | 44 | | | | | | | | |
| AC-11m | | | | | | | | | | | | 13 | | | | | | | | | | | | | | | | | | | | | | | | | | | | |
| AC-12m | | | | | | | | | | 5 | | 1 | | | | | | | | | | | | | | | | | | | | | | | | | | | | 13 |
| AC-13/34m | | | | | | | | | | | | | | | | | | | | | | | | | | | | | | | | | 1 | | | | | | | |
| AC-13m | | | | | | | 72 | | | | | | | | | | | | 1 | | | | 2 | | | | | | 1 | | | 1 | 1 | | | | | | | |
| AC-14m | | | | | | | | 106 | | | | | | | | | | | | | | | 1 | | | | | | 1 | | | | | | | | | | | |
| AC-15m | | | | 104 | | | | | | | | | | | | | | | | | | | | | | | | | | 1 | | 5 | | | | | | | | |
| AC-16m | | | 1 | | | | 1 | | | | | | | | | | | 70 | | | | | | | | | | | | | | | | | | | | | | |
| AC-17m | | 1 | | | | | | | | | | | | | | | 54 | | | | | | | 1 | | | | 1 | | | | | | | | | | | | |
| AC-18m | | | 88 | 1 | | | | | | | | | | | | | | | | | | | | | | | | | | | | | | | | | | | | |
| AC-19m | | | | | | | | | | | | | 1 | | | | | | | | | | | | | | | | | 1 | 79 | | | | | | | | | |
| AC-20a-m | | 11 | | | | | | | | | | | | | | | | | | | | | | | | | | | | | | | | | | | 24 | | | |
| AC-20a/b-m | | 1 | | | | | | | | | | | | | | | | | | | | | | | | | | | | | | | | | | | 1 | | | |
| AC-20b-m | | 69 | | | | | | 1 | | | | | | | | | | | | | | | | | | | 1 | | | | | | | | | | 5 | | | |
| AC-22m | | | | | | | | | | | | | | | | | | | | | | | 44 | | | | | | | | | | | | | | | | | |
| AC-23m | | | | 1 | | | | | | | | | | | 60 | 1 | | | | | | 2 | | | | | | | | | | | | | | | | | | |
| AC-24m | | | | | | | | | 3 | | | | | | | | | | | | | | | 1 | | | | | | | | | | | | | | | 33 | |
| AC-25m | | | | 1 | | | | | | | | | | | | | | | | | | | | | | | | | | | | | | | | 40 | | | | |
| AC-26m | | | | | | | | | | | | | | 102 | | | | | | | | | | | | | | | | | | | | | | | | | | |
| AC-27m | | | | | | | | | | 55 | | | | | | | | | | | | | | | | | | | | | | | | | | | | | | 4 |
| AC-28m | | | | | | | 1 | | | | | | | | | | | | | | | | | | | | | 22 | | | | | | | | | | | | |
| AC-29m | | | | | | | | | | | | | 60 | | | | | | | | | | | | | | | | | | | | | | | | | | | |
| AC-30m | | | | | | | | | | | | | | | | | | | 29 | | | | | | | | | | | | | | | | | | | | | |
| AC-31m | 1 | | | | | | | | | | | | | | | | | | 1 | | | | | | | 63 | | | | | | | | | | | | | | |
| AC-32m | | | | | | 1 | | | | | | | | | | | | | | | | | | 59 | | | | | | | | | | | | | | | | |
| AC-33m | | | 1 | | | | | | 1 | | | | | | | | | | | 96 | | | | 1 | | | | | | | | | | | | | | | | |
| AC-34m | | | | 1 | | | 2 | | 1 | | | | | | | | | | | | | | | | 1 | | | | | | 1 | | 16 | | | | | | | |
| AC-35m | | | | | 3 | | | | | | | | | | | | | | | | | | | | | | | | | | | | | | 57 | | | | | |
| AC-36m | 1 | | | | | | | | | | | | | 1 | | 69 | | | | | | | | | | | | | | 1 | | | | | | | | | | |
| AC-37m | | | | | | | | | | | | | 1 | | | | | | | | | | | | | | | | | | | 1 | 1 | 35 | | | | | | |

**B**

| | 1 | 2 | 3 | 4 | 5 | 6 | 7 | 8 | 9 | 10 | 11 | 12 | 13 | 14 | 15 | 16 | 17 | 18 | 19 | 20 | 21 | 22 | 23 | 24 | 25 | 26 | 27 | 28 | 29 | 30 | 31 | 32 | 33 | 34 | 35 | 36 | 37 | 38 | 39 | 40 |
|---|---|---|---|---|---|---|---|---|---|---|---|---|---|---|---|---|---|---|---|---|---|---|---|---|---|---|---|---|---|---|---|---|---|---|---|---|---|---|---|---|
| AC-1+21f | 1 | | | | 1 | | | | | | 98 | | | | | 1 | | | | | | | | | 23 | | | | | | 1 | | | | | | | 27 | | |
| AC-2f | | 1 | | | | | | | | | | | | | | | | | | | 79 | | | | | | | | | | | | 1 | | | | | | | |
| AC-3f | 1 | 1 | | 2 | | | | | 39 | | | | | 1 | | | | | | | | | | | | | | | | | | | | | | | | | 3 | |
| AC-4p-f | | | | | | | | | | | | | | | | | | 1 | | | | | | | | | | | | 40 | | | | | 1 | | | | | |
| AC-4q-f | 156 | | 1 | | 1 | | | | | | 1 | | | | | | | | | | | | | | | 1 | | | | 1 | | | | | | | | | | |
| AC-5f | | | | | | 1 | | | | | 1 | | | 1 | | | | | | | | 97 | | | | | | | | | | | | | | | | | | |
| AC-6f | | | | 2 | 114 | | | | | | | | | | | | | 1 | | | 1 | | | | 1 | | | | | | | | | | 3 | 1 | | | | |
| AC-7f | | | | | | | | | | | | | | | | | | | | | | | | | | | | | 49 | | | | | | | | | | | |
| AC-8f | | | | | | 66 | | | | | | | | | | | | | | | | | | | | | | | | | | | | | | | | | | |
| AC-9f | 2 | | | 2 | | | | 1 | | | | | | | | | | | | | | | | | | | 76 | | | | | | | | | | | | | |
| AC-10f | | | | | | | | | | | | | 1 | | | | | | | | | | | | | | | | | | | 24 | | | | | | | | |
| AC-11f | | | 1 | | | | | | | | | 15 | | | | | | | | | | | | | | | | | | | | | | | | | | | | |
| AC-12f | 1 | | | | | | | | | 3 | | | | | | | | | | | | | | | | | | | | | | | | 1 | | | | | | 22 |
| AC-13/34f | | | | | | | | | | | | | | | | | | | | | | | | | | | | | | | | | 1 | | | | | | | |
| AC-13f | | | | | | | 60 | | | | | | | 1 | | | | | | | 1 | | | | | | | | | | | | 2 | | | | | | | |
| AC-14f | | | | | | | | 123 | | | | | | | | | | | | | | | 1 | | | | | | 1 | | | | | | | | | | | |
| AC-15f | | | | 101 | | | | | | | | | | | | | | | | 1 | | 1 | | | | | | | | | | 5 | | | | | | | | |
| AC-16f | | | 1 | | | | | | | | | | | | | | | 86 | | | | | | | | | | | | | | | | | | | | | | |
| AC-17f | | | | | | 1 | | | | | | | | | | | 50 | | | | | | | | | | | | | | | | | | 1 | | | | | |
| AC-18f | | | 87 | | | | | | | | | | | | | | | | | | | | | | | | | | | | | | | | | 3 | 1 | | | |
| AC-19f | | | | | | | | | | | | | 1 | | 1 | | | | | | | | | | | | | | | 1 | 86 | | | | | | | | | |
| AC-20a-f | | 10 | | | 1 | | | | | | | | 1 | | | | | | | | | | | | | | | | | | | | | | | | 21 | | | |
| AC-20b-f | | 140 | | | | | | 1 | | | | | | | | | | | | | | | | | | | 1 | | 1 | | | | | | | | 3 | | | |
| AC-22f | | | | | | | | | | | | | | | | | | | | | | | 46 | | | | | | | | | | | | | | | | | |
| AC-23f | | | | 1 | | | | | | | | | | | 53 | | | | | | | 2 | | | | | | | | | | | | | | | | | | |
| AC-24f | | | | | | | | | 8 | | | | | | | | | | | | | 1 | | | | | | | | | | | | | | | | | 26 | |
| AC-25f | | | | 1 | | | | | | | | | | | | | | | | | | | | | | | | | | | | | | | | 39 | | | | |
| AC-26f | | | | | | | | | | | | | | 93 | | | | | | | | | | | | 1 | | | | | | | | | | | | | | |
| AC-27f | | | | | | | | | | 82 | | | | | | | | | | | | | | | | | | | | | | | | | | | | | | 6 |
| AC-28f | 1 | | | | | | | | | | | | | | | | | | | | | | | 1 | | | | 49 | | | | | | | | | | | | |
| AC-29f | | | | | | | | | | | | | 49 | | | | | | | | | | | | | | | | | | | | | | | | | | | |
| AC-30f | | | | | | | 1 | | | | | | | | | | | | 28 | | | | | | | | | | | | | | | | | | | | | |
| AC-31f | 1 | | | | | | | | | | | | | | | | | | | | | | | | | 71 | | | 1 | | | | | | | | | | | |
| AC-32f | | | | 1 | | | | | | | | | | | | | | | | | | | | 89 | | | | | | | | | | | | | | | | |
| AC-33f | | | 1 | | | | | | 1 | | | | | | | | | | | 97 | | | | | | | | | | | | | | | | | | | | |
| AC-34f | | | 1 | 1 | | | 1 | | | | | | | | | | | | | | | | | | | | | | | | | | 19 | | | | | | | |
| AC-35f | | | | | 6 | | | | | | | | | | | | | | | | | | | | | | | | | | | | | | 42 | | | | | |
| AC-36f | 1 | | | | | | | | | | | | | | | 55 | | | | | | | | | | | | | | 1 | 1 | | | | | | | | | |
| AC-37f | | | | | | | | | | | | | | | | | | | | | | | | | | | | | | | | | 1 | 58 | | | | | | |


1. Nugent, C. M., Easton, A. A., Norman, J. D., Ferguson, M. M. & Danzmann, R. G. A SNP based linkage map of the arctic charr (Salvelinus alpinus) genome provides insights into the diploidization process after whole genome duplication. *G3: Genes, Genomes, Genetics* **7**, 543–556 (2017).

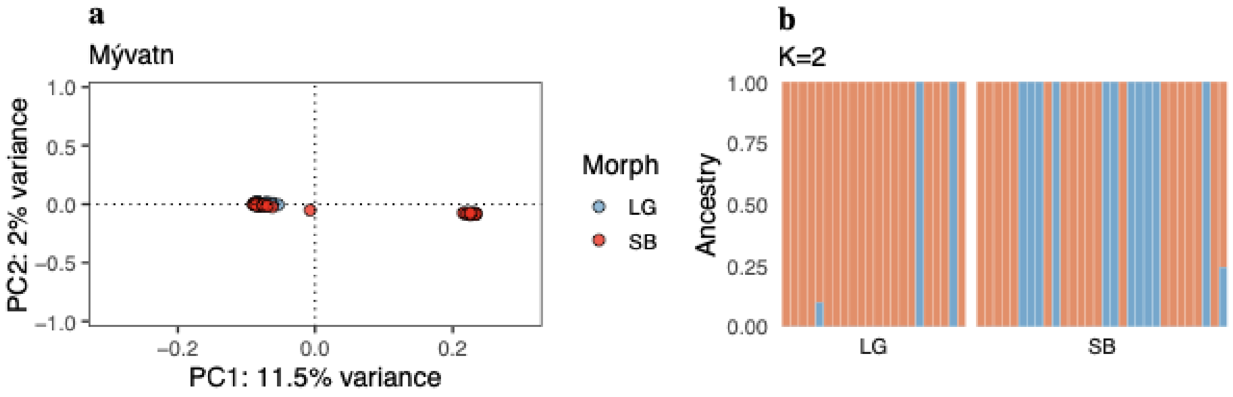


**Supplementary Fig. 2 Genetic differentiation among Arctic charr morphs in Lake Mývatn**. **(a)** Scores of individuals along PC1 and PC2, and **(b)** ancestry proportions of individuals assuming two clusters (K=2), estimated based on a downsampled set of 0.51 million SNPs (MAF > 0.05). The pre-assignment of morphs was based on sampling habitats. Morph abbreviations: Large generalist (LG) and Small benthic (Krús, SB).

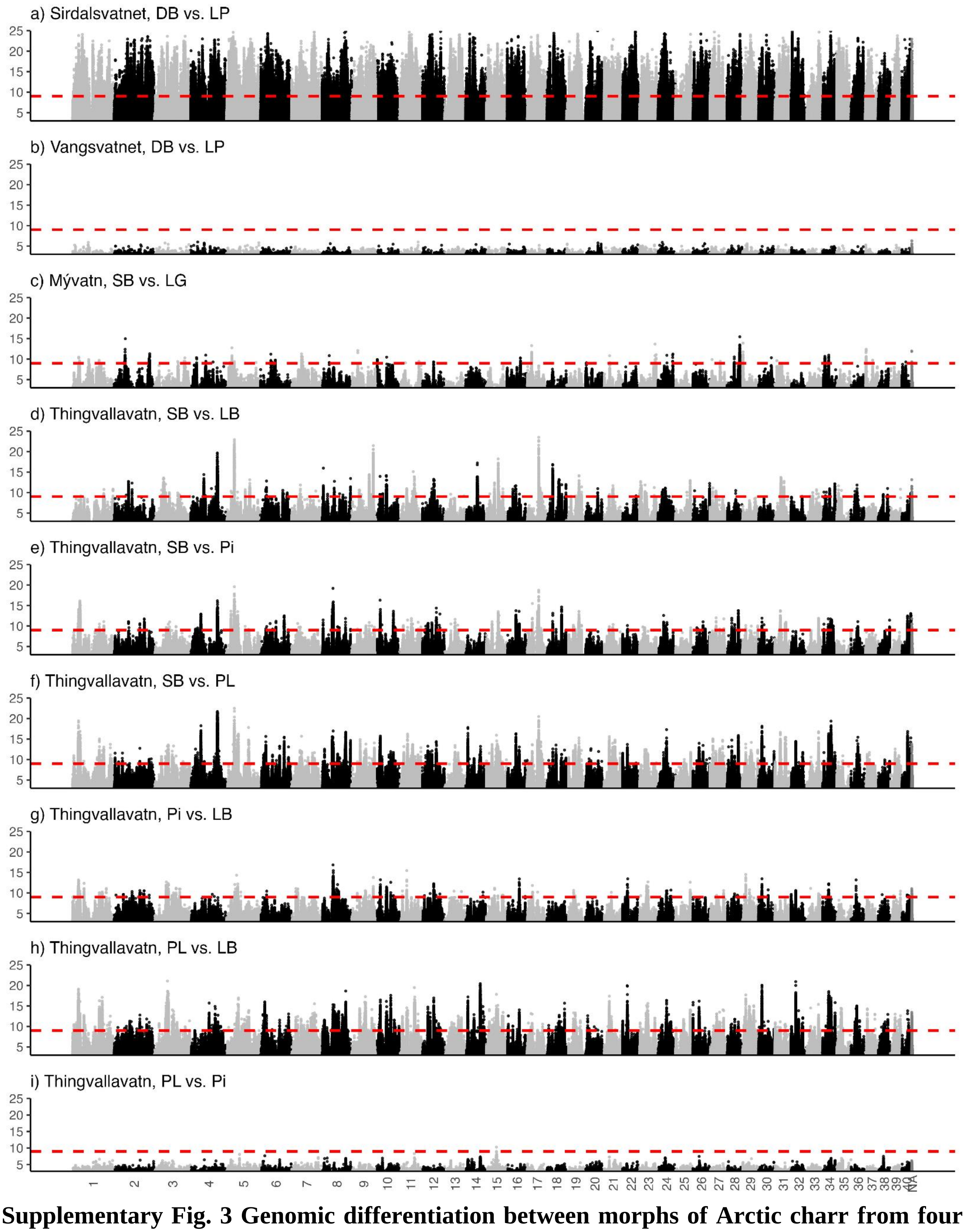


**Supplementary Fig. 3 Genomic differentiation between morphs of Arctic charr from four lakes.** Genome scan based on estimated allele frequencies for individual SNPs. (**a)** Sirdalsvatnet, (**b)** Vangsvatnet, (**c)** Mývatn, and (**d–i)** Thingvallavatn. The x-axis represents the scaffolds, and the y-axis represents the significance value (-log10(*P*-value) per SNP. Each dot corresponds to a single SNP. The horizontal red line indicates the significance threshold based on Bonferroni correction with the adjusted significance level for $\alpha = 10^{-3}$. Morph abbreviations for Sirdalsvatnet and Vangsvatnet: Dwarf benthic (DB) and Large pelagic (LP); for Mývatn, Large generalist (LG) and Small benthic (Krús, SB), and Thingvallavatn, Piscivorous (Pi), Planktivorous (PL), Large benthivorous (LB) and Small benthivorous (SB).

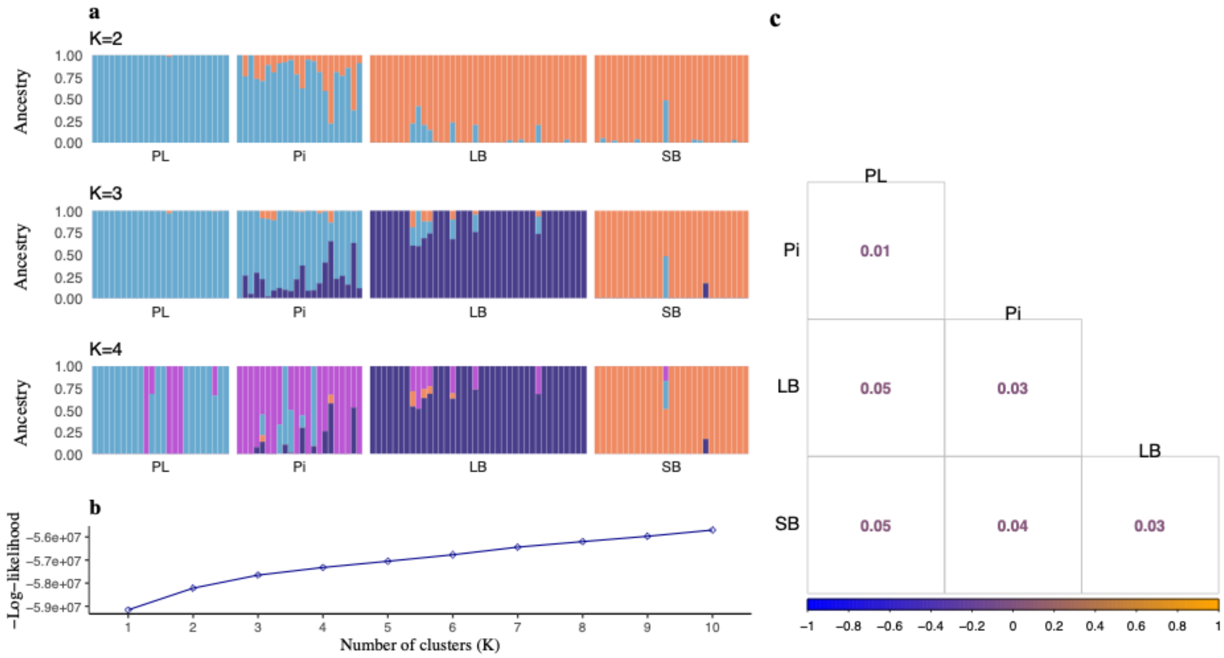


**Supplementary Fig. 4 Admixture analysis of individuals from Lake Thingvallavatn. (a)** Ancestry proportions generated in NGSAdmix based on a downsampled set of 0.58 million SNPs (MAF > 0.05). Morph abbreviations: Piscivorous (Pi), Planktivorous (PL), Large benthivorous (LB), and Small benthivorous (SB). (**b)** Log-likelihoods for 1–10 clusters in the admixture analysis. K = 3 was determined to best represent the genetic structure in the lake, aligning with the three distinct groups observed in the PCA plot (Fig. 3g**)**. Increasing K beyond 3 resulted in only minor changes in admixture proportions, suggesting that three main genetic groups capture the primary population structure within this lake.

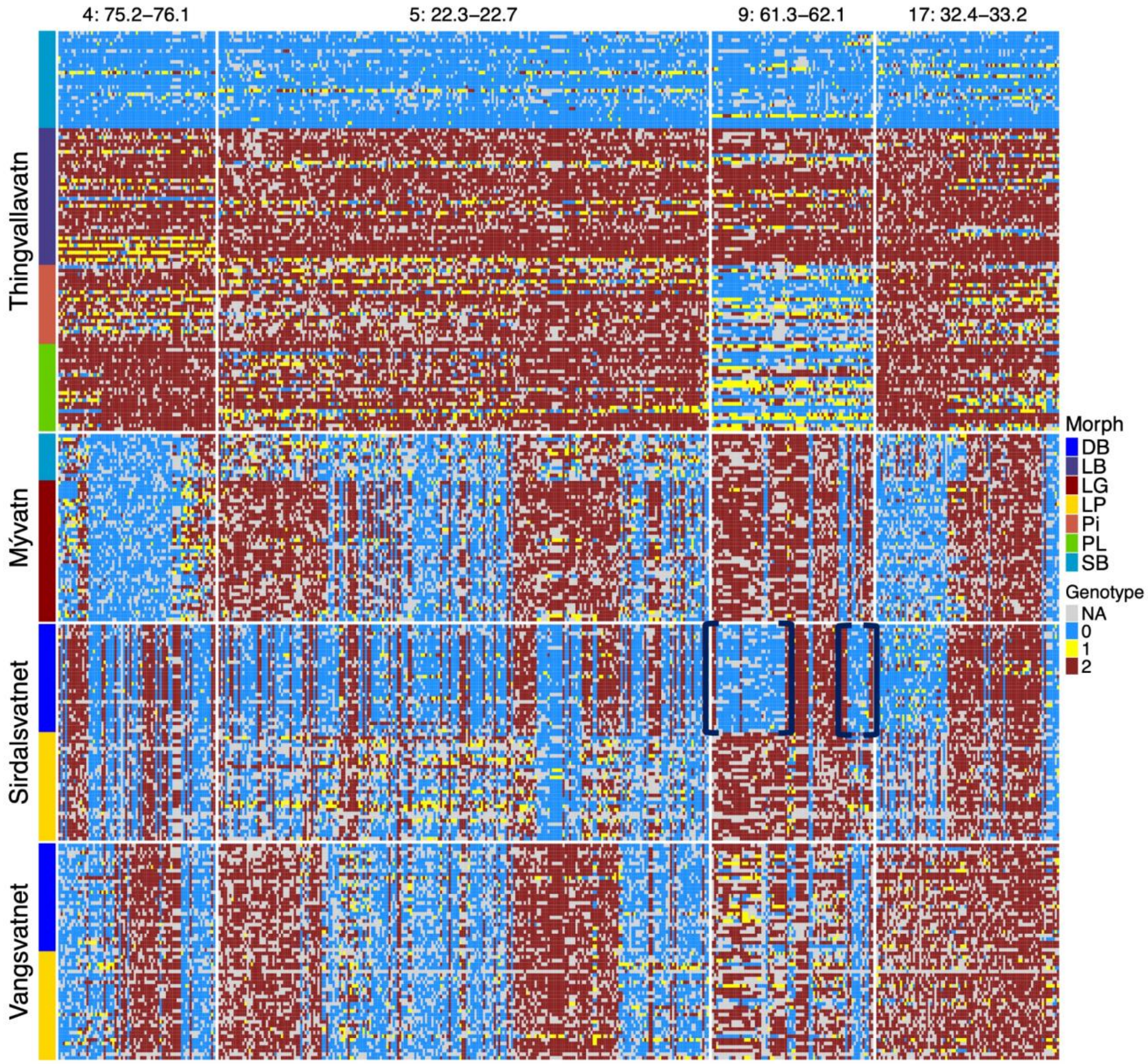


**Supplementary Fig. 5 Predicted genotypes based on genotype likelihoods from diagnostic markers at four putative inversions distinguishing large and small benthic morphs in Thingvallavatn.** Genotype distributions sorted by lake and morph are shown for the putative inversion regions on scaffolds 4, 5, 9, and 17. Morph abbreviations for Sirdalsvatnet and Vangsvatnet: Dwarf benthic (DB) and Large pelagic (LP); for Mývatn, Large generalist (LG) and Small benthic (Krús, SB), and Thingvallavatn, Piscivorous (Pi), Planktivorous (PL), Large benthivorous (LB) and Small benthivorous (SB). Individuals are colored according to their estimated genotype. The black brackets indicate a shared pattern of differentiation between morphs.

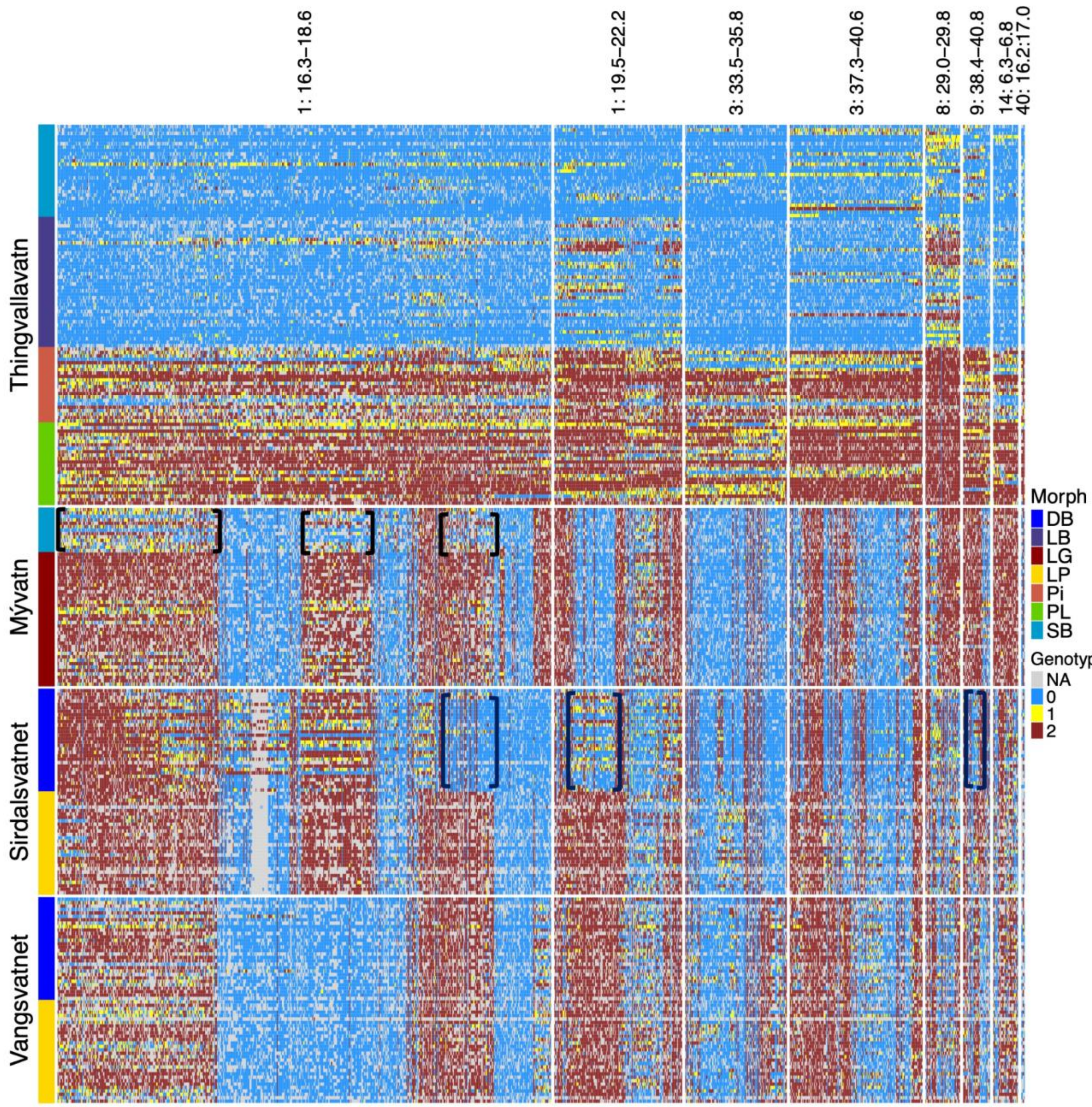


**Supplementary Fig. 6 Predicted genotypes based on genotype likelihoods from diagnostic markers at six putative inversions distinguishing Arctic charr benthic (large and small benthivorous) and pelagic (piscivorous and planktivorous) morphs in Thingvallavatn.** Genotype distributions sorted by lake and morph are shown for the putative inversion regions on scaffolds 1, 3, 8, 9, 14, and 40. Morph abbreviations for Sirdalsvatnet and Vangsvatnet: Dwarf benthic (DB) and Large pelagic (LP); for Mývatn, Large generalist (LG) and Small benthic (Krús, SB), and Thingvallavatn, Piscivorous (Pi), Planktivorous (PL), Large benthivorous (LB) and Small benthivorous (SB). Individuals are colored according to their estimated genotype. The black brackets indicate a shared pattern of differentiation between morphs.

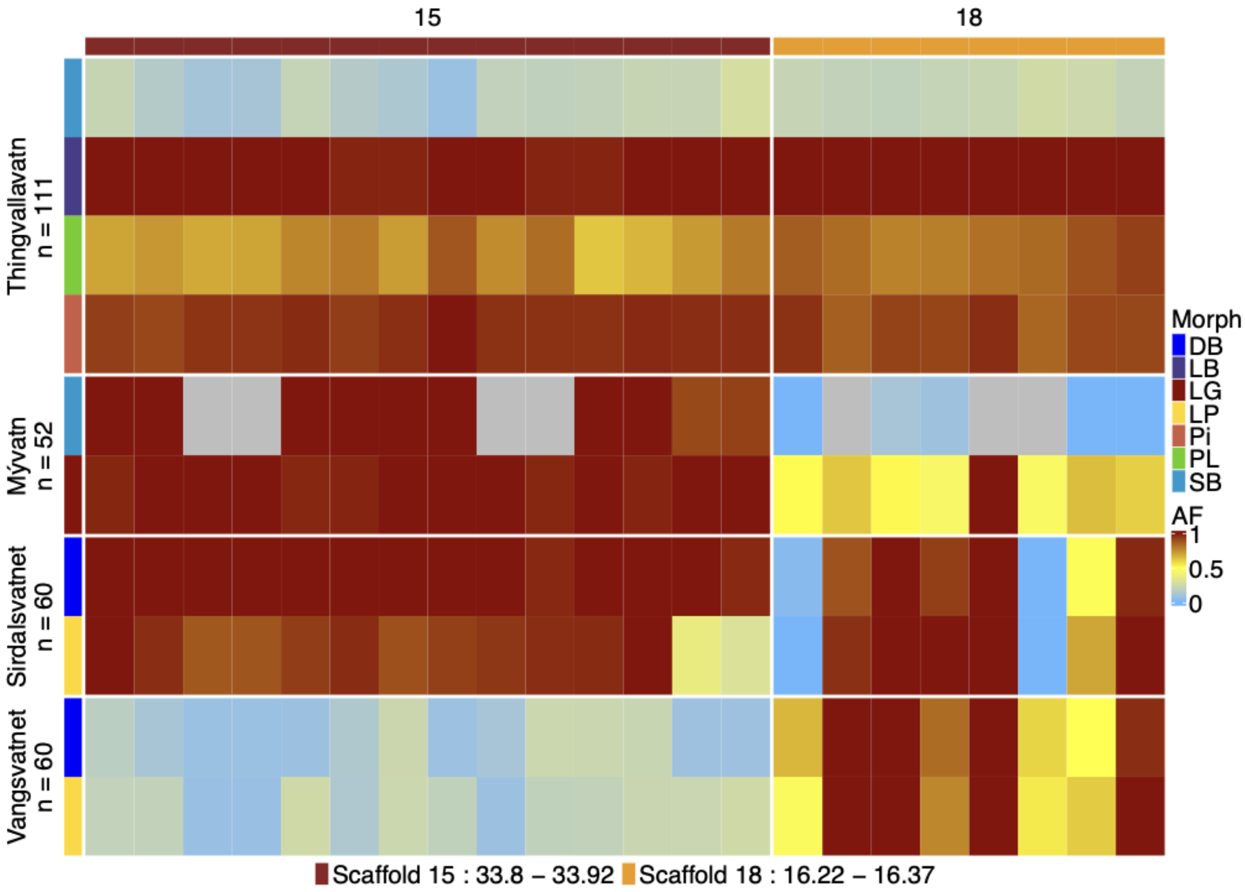


**Supplementary Fig. 7 Allele frequency of diagnostic SNPs outside the putative inversions selected based on the small/large benthivorous contrast of Arctic charr morphs from Thingvallavatn.** Each column title denotes a scaffold, with each value representing a diagnostic marker. The colors of the column annotation panel denote haplotype. Haplotypes were defined based on a windowed approach, where the maximum distance allowed between adjacent single nucleotide polymorphisms (SNPs) within the same haplotype was set to 50 kilobases (kb). This threshold was chosen to capture localized linkage disequilibrium patterns, ensuring that SNPs within each haplotype block were tightly linked and likely to be inherited together. Only regions containing more than four SNPs were included to ensure sufficient density for linkage analysis. Each of the presented regions ranges in size from 0.116 to 0.146 Mb. SNPs were tracked based on the most common allele in the Small benthivorous morph from Thingvallavatn. Allele frequencies (AF) of these alleles are indicated by the color code, grey color indicates missing data. Morph abbreviations for Sirdalsvatnet and Vangsvatnet: Dwarf benthic (DB) and Large pelagic (LP); for Mývatn, Large generalist (LG) and Small benthic (Krús, SB), and Thingvallavatn, Piscivorous (Pi), Planktivorous (PL), Large benthivorous (LB) and Small benthivorous (SB).

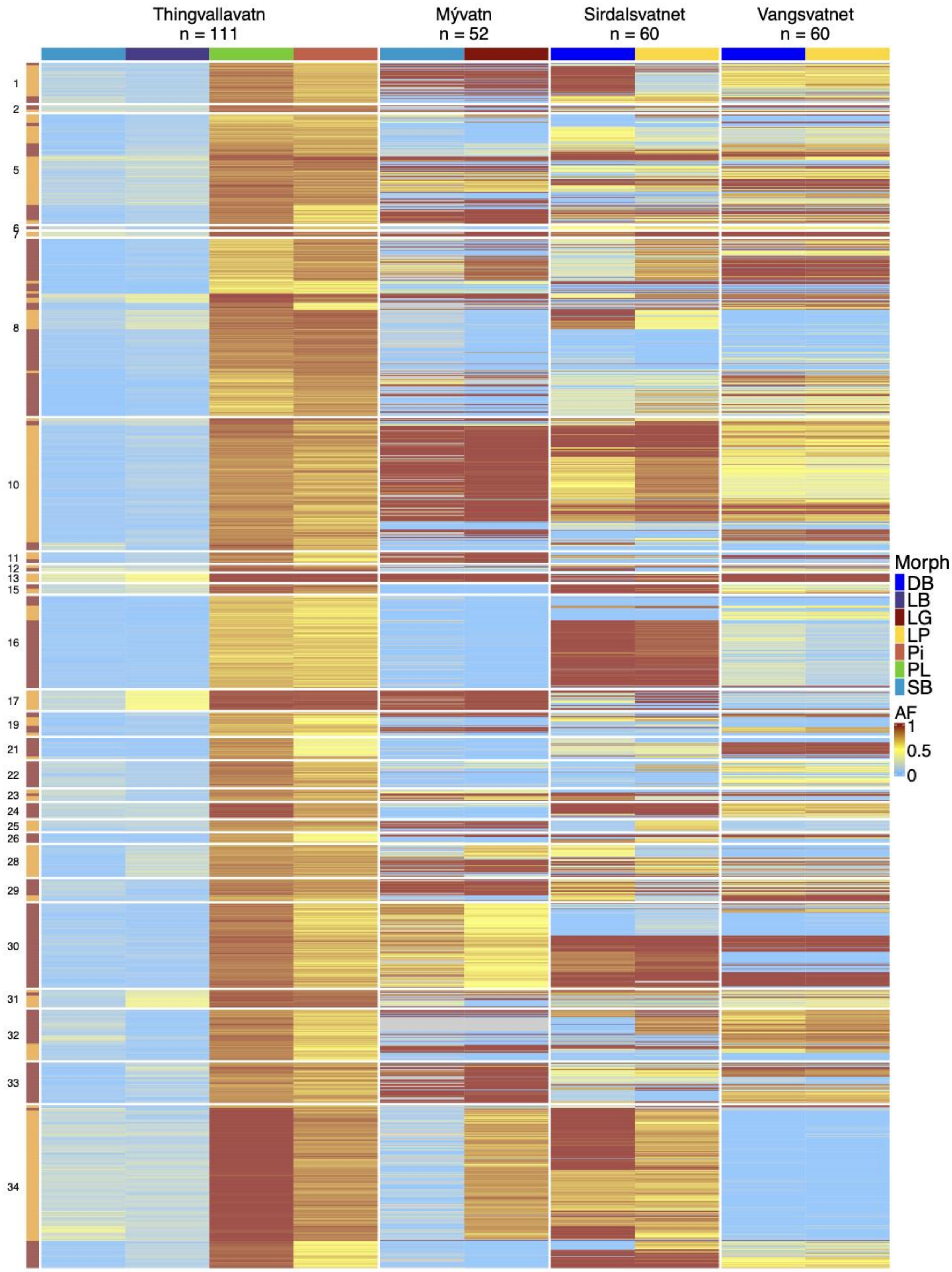

Thingvallavatn
n = 111
Mývatn
n = 52
Sirdalsvatnet
n = 60
Vangsvatnet
n = 60
1
2
5
6
7
8
10
11
12
13
15
16
17
19
21
22
23
24
25
26
28
29
30
31
32
33
34
Morph
DB
LB
LG
LP
Pi
PL
SB
AF
1
0.5
0

**Supplementary Fig. 8 Allele frequency of diagnostic SNPs outside the putative inversions selected based on the benthic/pelagic contrast of Arctic charr morphs from Lake Thingvallavatn.** Each row title denotes a scaffold, with each value representing a diagnostic marker. The colors on the left annotation panel denote haplotype. Haplotypes were defined as described in Supplementary Fig. 6. Each of the presented regions ranges in size from 0.167 to 0.314 Mb. SNPs were tracked based on the most common allele in the benthic morph from Thingvallavatn. Allele frequencies (AF) of these alleles are indicated by the color code, grey color indicates missing data. Morph abbreviations as in Supplementary Fig. 6.

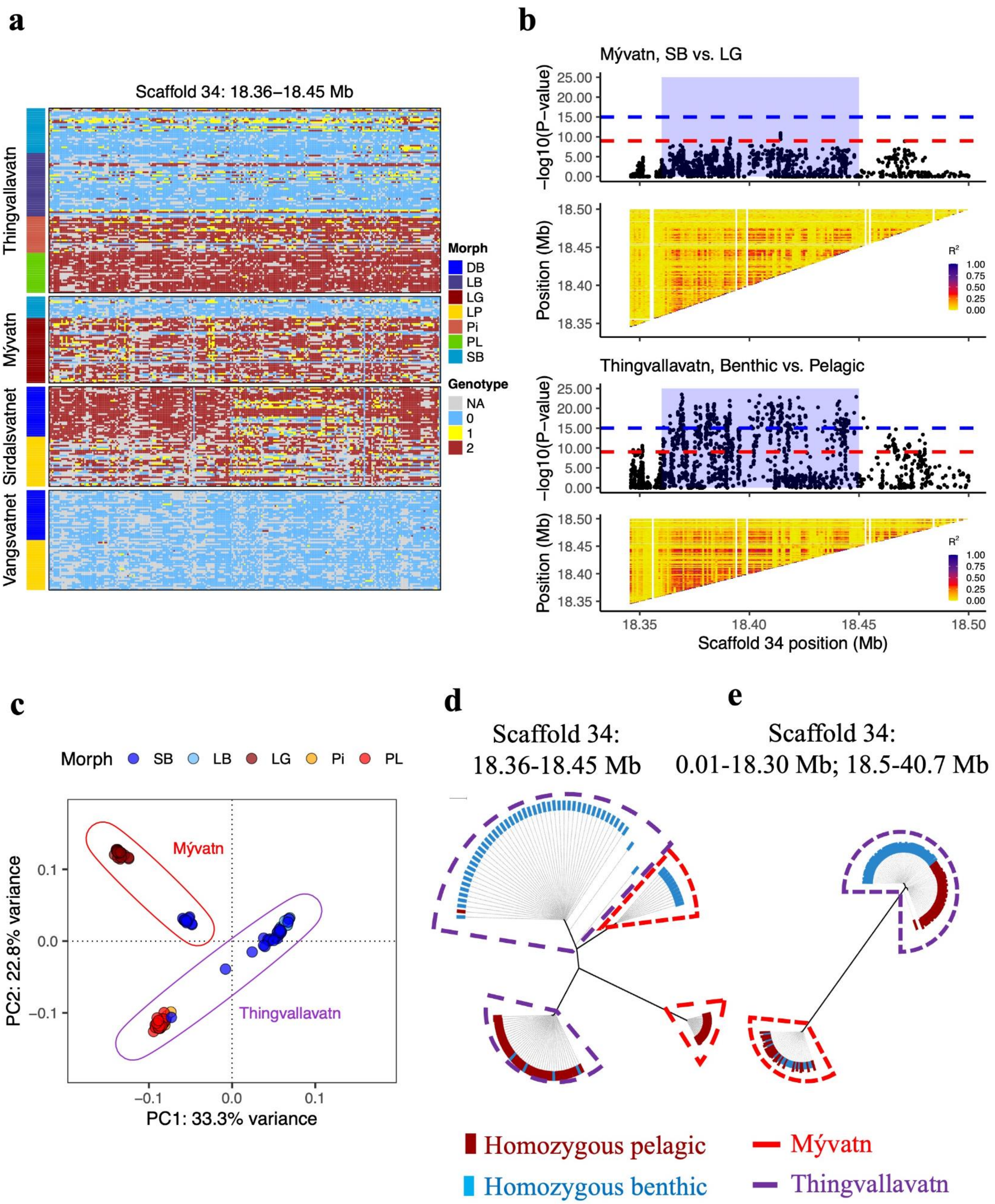


**Supplementary Fig. 9 Genetic differentiation at a region on scaffold 34 among Arctic charr morphs in Mývatn and Thingvallavatn. (a)** Predicted genotypes based on genotype likelihoods from diagnostic markers at this region sorted by lake and morph. Morph abbreviations as in Supplementary Fig. 6. Individuals are colored according to their estimated genotype. Each value in the column represents a diagnostic marker. SNPs were tracked based on the most common allele in benthic morphs from Thingvallavatn. **(b)** Zoom-in profile of the genome-wide scan based on estimated allele frequencies for individual SNPs and linkage disequilibrium represented as R2 among genotypes in the diagnostic region on scaffold 34 (18.36-18.45 Mb). **(c)** PCA plot showing individual clustering of samples homozygous for benthic or pelagic haplotype from Mývatn and Thingvallavatn within the diagnostic region (scaffold 34: 18.36-18.45 Mb). **(d)** Neighbor-joining tree of samples homozygous for benthic or pelagic haplotype from Mývatn and Thingvallavatn lakes within the diagnostic region (scaffold 34: 18.36-18.45 Mb) and **(e)** outside the diagnostic region (scaffold 34: 0.01-18.3 Mb and 18.5 - 40.7 Mb).

**Supplementary Table 1** Sample metadata, geological context, and history of the studied lakes where distinct Arctic charr morphs occur, including spawning times of the study populations. The Norwegian samples were collected from 1980 to 1984 and were the same as those used by Hindar et al. (1986) and they are maintained in a frozen tissue bank kept by L.L and N.R. at the Department of Zoology, Stockholm University. The samples from Thingvallavatn were collected in 2016-2018. The samples of LG-charr from Mývatn were taken in 2014 from a fisheries survey by the Freshwater Research Institute, and the samples of SB (Krús) charr were collected in 2015.

| Country | Location | Age in years, post-glacial period | Max depth, m | Morph | Abbreviated name | Lake zone | Sample size | Sample reference | Spawning |
|---|---|---|---|---|---|---|---|---|---|
| Norway | Vangsvatnet | ~10,000 | 60 | Large pelagic | LP | Pelagic | 30 | (Hindar et al. 1986) | late October-early December, shallow water (Jonsson & Hindar, 1982) |
| Norway | Vangsvatnet | - | - | Dwarf benthic | DB | Benthic | 30 | (Hindar et al.1986) | late October-early December, shallow water (Jonsson & Hindar, 1982) |
| Norway | Sirdalsvatnet | ~10,000 | 165 | Large pelagic | LP | Pelagic | 30 | (Hindar et al.1986) | late October -November, 0-32 m (Jonsson & Jonsson, 2001) |
| Norway | Sirdalsvatnet | - | - | Dwarf benthic | DB | Benthic | 30 | (Hindar et al., 1986) | throughout the year, peaks June-September, 55-70 m (Jonsson & Jonsson, 2001) |
| Iceland | Thingvallavatn | ~10 000 (Saemundsson, 1992) | 114 | Large benthivorous | LB | Benthic | 38 | Not published | Peak in late July - early August. Stony littoral zone near cold springs (Skúlason et al., 1989) |
| Iceland | Thingvallavatn | - | - | Small benthivorous | SB | Benthic | 27 | Not published | Variable, early September – November. Stony littoral zone. (Skúlason et al., 1989) |
| Iceland | Thingvallavatn | - | - | Planktivorous | PL | Pelagic | 24 | Not published | Peak in late September – mid-Octobe. Littoral zone (Skúlason et al., 1989) |
| Iceland | Thingvallavatn | - | - | Piscivorous | Pi | Benthic *Nitella* zone | 22 | Not published | Mid-September - mid-October. Littoral zone (Skúlason et al., 1989) |
| Iceland | Mývatn | ~2,300 (Einarsson et al., 2004) | 4.5 | Large generalist | LG | Pelagic/ Benthic | 22* | Not published | late October, peaks in November (Guðni Guðbergsson (personal communication)) |
| Iceland | Mývatn | - | - | Small benthic (Krús) | SB | Benthic | 30* | Not published | October, November, up to December (Guðni Guðbergsson (personal communication)) |

* Preliminary assignment

**Supplementary Table 2** Number of scaffolds and SNPs where at least one marker exceeds the Bonferroni corrected significance level ($\alpha = 10^{-3}$) identified per each contrast between Arctic charr morphs from the four lakes. Morph abbreviations for Sirdalsvatnet and Vangsvatnet: Dwarf benthic (DB) and Large pelagic (LP); for Mývatn, Large generalist (LG) and Small benthic (Krús, SB), and Thingvallavatn, Piscivorous (Pi), Planktivorous (PL), Large benthivorous (LB) and Small benthivorous (SB).

| Lake | Contrast | Number of scaffolds | Number of SNPs | Number of SNPs on unplaced scaffolds |
|---|---|---|---|---|
| Sirdalsvatnet | DB vs. LP | 40 | 103,326 | 1169 |
| Vangsvatnet | DB vs. LP | 0 | 0 | 0 |
| Mývatn | SB vs. LG | 20 | 198 | 2 |
| Thingvallavatn | SB vs. LB | 33 | 2,898 | 9 |
| Thingvallavatn | SB vs. PL | 39 | 9,122 | 31 |
| Thingvallavatn | SB vs. Pi | 32 | 2,377 | 4 |
| Thingvallavatn | LB vs. PL | 40 | 9,861 | 120 |
| Thingvallavatn | LB vs Pi | 30 | 563 | 5 |
| Thingvallavatn | PL vs. Pi | 1 | 1 | 0 |
| Thingvallavatn | Benthic vs. Pelagic | 40 | 24,852 | 142 |

**Supplementary Table 3** Genotype distribution at four putative inversions showing genetic differentiation between the small and large benthivorous and large benthic morphs present in Thingvallavatn.

| Group | Scaffold and region (start-end, Mb) | | | | | | | |
|---|---|---|---|---|---|---|---|---|
| | 4: 75.25-76.13 | | 5: 22.30-22.75 | | 9: 61.30-62.11 | | 17: 32.45-33.20 | |
| | SB | LB | SB | LB | SB | LB | SB | LB |
| Homozygous major (small benthivorous) | 26 | 1 | 26 | 0 | 26 | 1 | 24 | 0 |
| Heterozygous | 1 | 9 | 1 | 5 | 1 | 6 | 3 | 5 |
| Homozygous minor (large benthivorous) | 0 | 28 | 0 | 33 | 0 | 31 | 0 | 33 |
| Total | 27 | 38 | 27 | 38 | 27 | 38 | 27 | 38 |

**Supplementary Table 4** Nucleotide diversity (θ) assessed across the whole genome and at putative inversion regions among Arctic charr morphs from Thingvallavatn homozygous for small and large benthivorous haplotype

| Scaffold: start - end | Haplotype | Mean θ |
|---|---|---|
| Genome-wide | Small benthivorous | 0.0015 |
| Genome-wide | Large benthivorous | 0.0014 |
| 4: 75.25-76.13 | Small benthivorous | 0.0012 |
| | Large benthivorous | 0.0006 |
| 5: 22.30-22.75 | Small benthivorous | 0.0013 |
| | Large benthivorous | 0.0004 |
| 9: 61.30-62.11 | Small benthivorous | 0.0028 |
| | Large benthivorous | 0.0014 |
| 17: 32.45-33.20 | Small benthivorous | 0.0008 |
| | Large benthivorous | 0.0005 |

**Supplementary Table 5** Genotype distribution at eight putative inversions showing genetic differentiation between the benthic (B) and pelagic (P) morphs present in Thingvallavatn.

| | Scaffold and region (start-end, Mb) | | | | | | | | | | | | | | | |
|---|---|---|---|---|---|---|---|---|---|---|---|---|---|---|---|---|
| | 1: 16.30-18.60 | | 1: 19.50-22.20 | | 3: 33.50-35.80 | | 3: 37.35-40.60 | | 8: 29.05-29.83 | | 9: 38.40-40.80 | | 14: 6.33-6.87 | | 40: 16.25-17.01 | |
| Group | B | P | B | P | B | P | B | P | B | P | B | P | B | P | B | P |
| Homozygous major (benthic) | 62 | 5 | 31 | 0 | 62 | 6 | 52 | 3 | 22 | 0 | 48 | 2 | 56 | 8 | 26 | 1 |
| Heterozygous | 2 | 21 | 29 | 16 | 2 | 25 | 10 | 15 | 30 | 1 | 15 | 17 | 8 | 17 | 30 | 9 |
| Homozygous minor (pelagic) | 0 | 20 | 4 | 30 | 0 | 15 | 2 | 28 | 12 | 45 | 1 | 27 | 0 | 21 | 8 | 36 |
| Total | 64 | 46 | 64 | 46 | 64 | 46 | 64 | 46 | 64 | 46 | 64 | 46 | 64 | 46 | 64 | 46 |

**Supplementary Table 6** Nucleotide diversity (θ) assessed across the whole genome and at putative inversion regions among Arctic charr morphs from Lake Thingvallavatn homozygous for benthic or pelagic haplotype.

| Scaffold: start - end (Mb) | Haplotype | Mean pairwise theta |
|---|---|---|
| Genome-wide | Benthic | 0.0014 |
| Genome-wide | Pelagic | 0.0015 |
| 1: 16.30-18.60 | Benthic | 0.0018 |
| | Pelagic | 0.0025 |
| 1: 19.50-22.20 | Benthic | 0.0005 |
| | Pelagic | 0.0012 |
| 3: 33.50-35.80 | Benthic | 0.0006 |
| | Pelagic | 0.0010 |
| 3: 37.35-40.60 | Benthic | 0.0005 |
| | Pelagic | 0.0006 |
| 8: 29.05-29.83 | Benthic | 0.0008 |
| | Pelagic | 0.0015 |
| 9: 38.40-40.80 | Benthic | 0.0007 |
| | Pelagic | 0.0009 |
| 14: 6.33-6.87 | Benthic | 0.0004 |
| | Pelagic | 0.0017 |
| 40: 16.25-17.01 | Benthic | 0.0032 |
| | Pelagic | 0.0005 |

**Supplementary Table 7** Nucleotide diversity (θ) assessed across the whole genome among Arctic charr morphs and populations. Morph abbreviations for Sirdalsvatnet and Vangsvatnet: Dwaf benthic (DB) and Large pelagic (LP); for Mývatn, Large generalist (LG) and Small benthic (Krús, SB), and Thingvallavatn, Piscivorous (Pi), Planktivorous (PL), Large benthivorous (LB) and Small benthivorous (SB).

| Country | Lake | Morph | Mean pairwise theta |
|---|---|---|---|
| **Across populations** | | | |
| Norway | Sirdalsvatnet | All | 0.0025 |
| Norway | Vangsvatnet | All | 0.0024 |
| Iceland | Thingvallavatn | All | 0.0015 |
| Iceland | Mývatn | All | 0.0026 |
| **Across morphs** | | | |
| Norway | Sirdalsvatnet | DB | 0.0023 |
| Norway | Sirdalsvatnet | LP | 0.0023 |
| Norway | Vangsvatnet | DB | 0.0023 |
| Norway | Vangsvatnet | LP | 0.0023 |
| Iceland | Mývatn | SB | 0.0019 |
| Iceland | Mývatn | LG | 0.0020 |
| Iceland | Thingvallavatn | PL | 0.0014 |
| Iceland | Thingvallavatn | Pi | 0.0015 |
| Iceland | Thingvallavatn | LB | 0.0014 |
| Iceland | Thingvallavatn | DB | 0.0014 |
| Iceland | Thingvallavatn | Benthic | 0.0014 |
| Iceland | Thingvallavatn | Pelagic | 0.0015 |

**Supplementary Table 8** Genotype distribution at locus 18.36-18.45 Mb of scaffold 34 showing genetic differentiation between morphs homozygous for benthic and pelagic haplotype present in Thingvallavatn and Mývatn.

| Group | Scaffold 34: 18.36-18.45 Mb | | | | | | Total |
|---|---|---|---|---|---|---|---|
| | Mývatn | | Thingvallavatn | | | | |
| | LG | SB | SB | LB | PL | Pi | |
| Homozygous major (benthic) | 2 | 10 | 18 | 28 | 0 | 1 | 59 |
| Heterozygous | 9 | 3 | 8 | 7 | 0 | 6 | 33 |
| Homozygous minor (pelagic) | 28 | 0 | 1 | 3 | 24 | 15 | 71 |
| Total | 39 | 13 | 27 | 38 | 24 | 22 | 163 |
